# Metabolic and behavioral effects of neurofibromin result from differential recruitment of MAPK and mTOR signaling

**DOI:** 10.1101/2025.07.25.666841

**Authors:** Valentina Botero, Jenifer Barrios, Anneke Knauss, Ethan Rosendahl, Kenneth J. Colodner, Seth M. Tomchik

## Abstract

Neurofibromatosis type 1 results from mutations in the *Neurofibromin 1* gene and its encoded neurofibromin protein. This condition produces multiple symptoms, including tumors, behavioral alterations, and metabolic changes. Molecularly, neurofibromin mutations affect Ras activity, influencing multiple downstream signaling pathways, including MAPK (Raf/MEK/ERK) and PI3K/Akt/mTOR signaling. This pleiotropy raises the question of which pathways could be targeted to treat the disease symptoms, and whether different phenotypes driven by neurofibromin mutations exhibit similar or diverging dependence on the signaling pathways downstream of Ras. To test this, we examined metabolic and behavioral alterations in the genetically tractable *Drosophila* neurofibromatosis type 1 model. *In vivo* genetic analysis revealed that behavioral effects of neurofibromin were mediated by MEK signaling, with no necessity for Akt. In contrast, metabolic effects of neurofibromin were mediated by coordinated actions of MEK/ERK and Akt/mTOR/S6K/4E-BP signaling. At the systemic level, neurofibromin dysregulated metabolism via molecular effects of Nf1 in interneurons and muscle. These changes were accompanied by altered muscle mitochondria morphology, with no concomitant changes in neuronal ultrastructure or neuronal mitochondria. Overall, this suggests that neurofibromin mutations affect multiple signaling cascades downstream of Ras, which differentially affect metabolic and behavioral neurofibromatosis type 1 phenotypes.

## Introduction

Neurofibromatosis type 1 (NF1) is a multisystemic genetic disorder characterized by a spectrum of clinical manifestations, including tumors such as cutaneous and plexiform neurofibromas, optic gliomas, and malignant peripheral nerve sheath tumors. Cutaneous symptoms include neurofibromas and a range of cutaneous, skeletal, and ocular symptoms. Additional manifestations include neurological symptoms and growth/metabolism-related symptoms such as short stature, metabolic changes, sleep disruption, cognitive/behavioral alterations, and fatigue [1–12]. The condition is caused by loss of function mutations in the Neurofibromin 1 gene, which encodes a large protein called neurofibromin (Nf1). Nf1 is a major Ras-GAP, which directly binds Ras, accelerates the hydrolysis of Ras-GTP, and thereby functions as a negative regulator of Ras signaling [13, 14]. Loss of Nf1 upregulates the activity of multiple downstream pathways, including mitogen-activated protein kinase (MAPK) signaling (Raf-MEK-ERK) and PI3K/Akt/mTOR. Further, cAMP/PKA levels are reduced [12, 15–20].

The MAPK signaling pathway, MEK in particular, is a target for treatment of plexiform neurofibromas in NF1 [21, 22], though with potentially significant adverse reactions [23–25]. This raises several questions, including whether other NF1 symptoms could be targeted with similar approaches. NF1 patients exhibit a range of nontumor symptoms, including increased frequency of learning deficits, attention-deficit/hyperactivity disorder, and autism spectrum disorder [26]. These cognitive and behavioral symptoms presumably result from neuronal dysfunction caused by the mutations in the *Neurofibromin 1* gene. Animal models exhibit deficiencies in behavioral tasks and neuronal function indicative of cognitive dysfunction [2, 12, 27–29]. These result from alterations in Ras/ERK and/or dopamine/cAMP signaling. The potential efficacy of targeting MEK (or other MAPK signaling molecules) for treating cognitive and behavioral symptoms in NF1 is unknown. A second major question is whether other signaling pathways could be targeted to treat NF1 symptoms – either alone or in combination with MEK – possibly with fewer side effects.

Metabolic alterations represent an understudied aspect of NF1 which may contribute to the development of other symptoms [11, 12, 30–32]. Patients experience a range of metabolic changes, including reduced cerebral glucose metabolism in the thalamus [33], lower blood glucose [34], reductions in muscle mass [35], decreased muscular strength and force [32, 36], alterations in muscle growth and metabolism, and increased resting energy expenditure [32]. Mouse models of NF1 similarly exhibit metabolic phenotypes, including altered muscle growth and muscle lipid mobilization [30, 37–39]. In the fruit fly, *Drosophila* melanogaster, loss of Nf1 alters metabolism [12], increasing energy expenditure [11], reducing triglyceride [11] and glycogen [40] levels, increasing fat turnover [11], increasing ROS production and decreasing lifespan [41], shifting locomotor activity [42–45], and altering metabolism-sleep interactions [40, 46]. Some of these effects are mediated by Nf1 in neurons [11, 40, 42–46], suggesting that central control of metabolism is affected. Though patient and animal model studies show metabolic dysfunction caused by the loss of Nf1 function, the underlying mechanisms are unclear. Furthermore, it is unknown whether the molecular mechanisms underlying metabolic alterations are similar to those driving cognitive and behavioral alterations.

Since there are multiple phenotypes in NF1, and the neurofibromin protein is a Ras-GAP that exerts pleiotropic effects on multiple signaling pathways downstream of Ras, a major question is whether different downstream signaling pathways exhibit differential involvement in NF1 phenotypes. For instance, are the NF1 cognitive/behavioral symptoms more dependent on one pathway (e.g., MEK) while the metabolic alterations more dependent on another (e.g., mTOR)? We tested this using the genetically tractable *Drosophila* NF1 model, which allows precise dissection of signaling pathway function *in vivo*. Using this model, we compared the signaling requirements for two NF1 phenotypes in *Drosophila*: behavioral alterations and metabolic dysregulation.

## Results

### Neurofibromin exerts behavioral effects via MEK, with no requirement for Akt signaling

To dissect the mechanisms by which Neurofibromin 1 affects organismal physiology, we first examined its effect on behavior in *Drosophila*. Genomic *nf1* mutations or panneuronal Nf1 loss of function increases spontaneous grooming in an Nf1 Ras GAP-related domain-dependent manner [43]. To test the underlying cellular signaling mechanisms downstream of Ras, we implemented an *in vivo* genetic analysis approach. Nf1 was knocked down pan-neuronally with RNAi [11, 43, 44], either alone or in combination with knockdown of a second signaling molecule downstream of n.

Ras (Fig. 1B). Flies were monitored in open field arenas for 5 minutes and grooming behavior was quantified (Fig. 1A,C). The phenotype is present in both males and females [45]. In the first set of experiments, we targeted the MAPK signaling cascade, knocking down Nf1 along with MEK (Fig. 1B,C). MEK is a known modifier of neurofibromatosis type 1 phenotypes and therapeutic target for treatment of plexiform neurofibromas [21, 22]. Knocking down Nf1 in neurons increased spontaneous grooming frequency (Fig. 1C), as expected [43–45]. Knockdown of MEK did not, on its own, affect behavior (Fig. 1C). However, when knocking down both MEK and Nf1, the MEK knockdown occluded the behavioral effect of knocking Nf1 down (Fig. 1C). This suggested that MEK was necessary for the Nf1 effect on the behavioral phenotype. To validate the double RNAi approach, we tested whether introducing a second RNAi alters Nf1 mRNA relative to the single Nf1 RNAi via quantitative PCR (qPCR). Nf1 mRNA levels were significantly reduced in both the single Nf1 knockdown and double Nf1+MEK knockdowns, and there was no significant difference in Nf1 mRNA expression between the single and double knockdowns (Fig. 1E). This demonstrated that there was no dilution of the Nf1 knockdown when expressing a second RNAi line using the same Gal4 driver. Next, we examined how knocking down Nf1 and/or MEK affected downstream ERK phosphorylation. Western blot analysis showed increased phosphorylated ERK (pERK) following Nf1 knock down in neurons without affecting total ERK (Fig. 1F). Knocking down Nf1 and MEK concurrently occluded the effect of Nf1 knockdown, normalizing pERK levels. This demonstrated that the double RNAi lines effectively modulated the targeted signaling pathways (MEK/ERK), providing further validation of the genetic approach.

**Figure 1:**
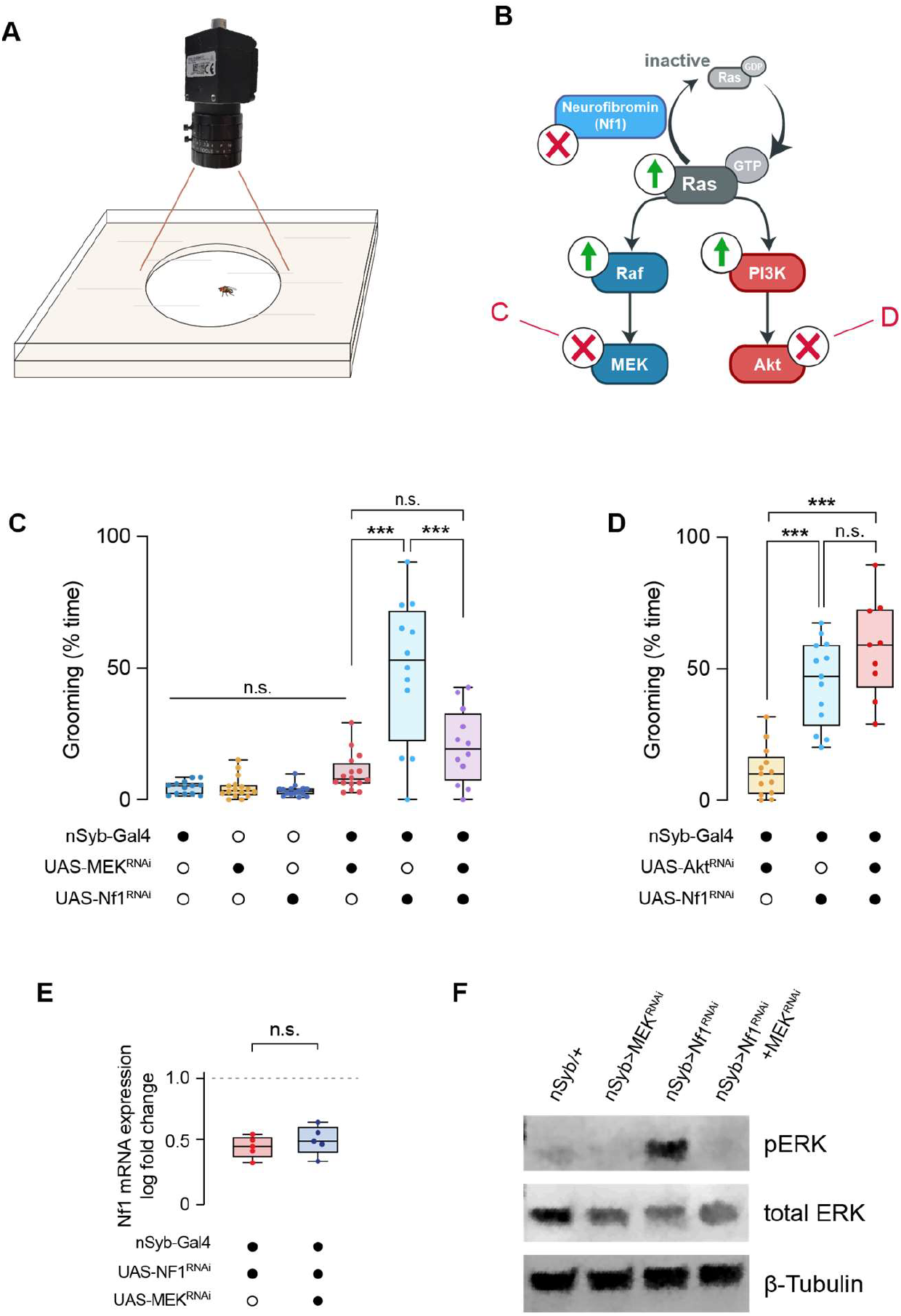
Loss of Nf1 alters a behavioral phenotype, grooming, via effects on MEK, but not Akt. **(A)** Diagram of the open field arena containing a fly, with a camera recording above. **(B)** Diagram showing Nf1-Ras signaling and the downstream Raf/MEK and PI3K/Akt signaling pathways. Green arrows represent the direction of signaling change following loss of Nf1. Red X marks show the molecules targeted for *in vivo* genetic analyses in panels C-D. **(C)** Quantification of grooming in flies harboring RNAi targeting Nf1, MEK, or both, along with controls. ***p < 0.001; n.s.: not significant (Šidák, n = 12-16). **(D)** Quantification of grooming in flies harboring RNAi targeting Nf1, Akt, or both. ***p < 0.001; n.s.: not significant (Šidák, n = 9-13). **(E)** Nf1 knockdown efficiency (log fold change normalized to Gal4/+ controls) in flies harboring RNAi targeting Nf1 or Nf1 + MEK, measured with quantitative PCR. n.s.: not significant (p = 0.42, Mann-Whitney, n = 5). **(F)** Representative western blot of total ERK and phosphorylated ERK in controls (nSyb-Gal4/+), and flies with pan-neuronal MEK knockdown, Nf1 knockdown, or Nf1 + MEK knockdown. Samples derived from the same experiment and processed in parallel, with β-tubulin as loading control. Representative blot shown from one of two experiments.

Nf1 modulates multiple signaling pathways downstream of Ras, including Raf/MEK and PI3K/Akt signaling pathways (Fig. 1B). Therefore, we next tested whether PI3K/Akt signaling is involved in regulating the behavioral phenotype. There are multiple PI3K isoforms [47], complicating loss-of-function analysis. Therefore, we tested Akt, which exists as a single isoform in *Drosophila* [48]. In contrast to MEK, knocking down Akt did not occlude the Nf1 effect on behavior – there was no significant difference in grooming between the single Nf1 knockdown and double Nf1 + Akt knockdown (Fig. 1D). This suggested that reducing the expression of MEK, but not Akt restored a behavioral phenotype (excessive grooming caused by loss of Nf1 in neurons). Thus, Nf1 affected behavior via selective effects on MAPK signaling in this signaling context.

### Metabolic effects of neurofibromin are mediated by coordinated actions of MEK/ERK and Akt/mTOR/S6K/4E-BP

The above data suggest Nf1 signals through MEK (but not Akt) to regulate behavior. Beyond behavioral regulation, Nf1 also regulates other phenotypes [11, 18, 20, 41, 46, 49–53]. Could these phenotypes exhibit differential dependence on Nf1-regulated signaling cascades? To begin exploring this, we examined Nf1-dependent metabolic regulation. Loss of Nf1 disrupts metabolic homeostasis, increasing CO_2_ production (and O_2_ consumption) [11, 12, 54], which correlates with organismal metabolic rate [55]. Like the behavioral phenotype [43], the Nf1-dependent metabolic phenotype is Ras dependent [11]. We utilized this phenotype to probe the cellular signaling mechanisms of Nf1 effects on metabolism (Fig. 2A). Prior metabolic studies focused on male flies, so we initially compared CO_2_ production between *nf1*^*P1*^ mutants and wCS10 controls in both sexes. In both males and females, there was a significant increase in CO_2_ production in *nf1*^*P1*^ mutants relative to controls (Fig. 2B). The Nf1 metabolic effect maps to a set of neurons labeled with the PCB-Gal4 driver [11]. When knocking Nf1 down in these neurons, CO_2_ production was increased (Fig. 2C). We knocked Nf1 down in these metabolism-regulating neurons to examine the dependence of the metabolic phenotype on signaling cascades downstream of Ras, using a similar double knockdown approach as above (Fig. 2D). First, we examined MAPK signaling, beginning with MEK. Knocking down MEK alone did not affect metabolic rate (CO_2_ production), but combining it with Nf1 knockdown occluded the effect of the Nf1 knockdown on metabolic rate (Fig. 2E). Moving one step downstream in the MAPK pathway, we observed a similar effect with ERK. Knocking down ERK alone did not affect metabolic rate, yet it occluded the Nf1 effect on metabolic rate when both were knocked down simultaneously (Fig. 2F). This suggested that MEK/ERK signaling is required for Nf1-dependent metabolic modulation.

**Figure 2:**
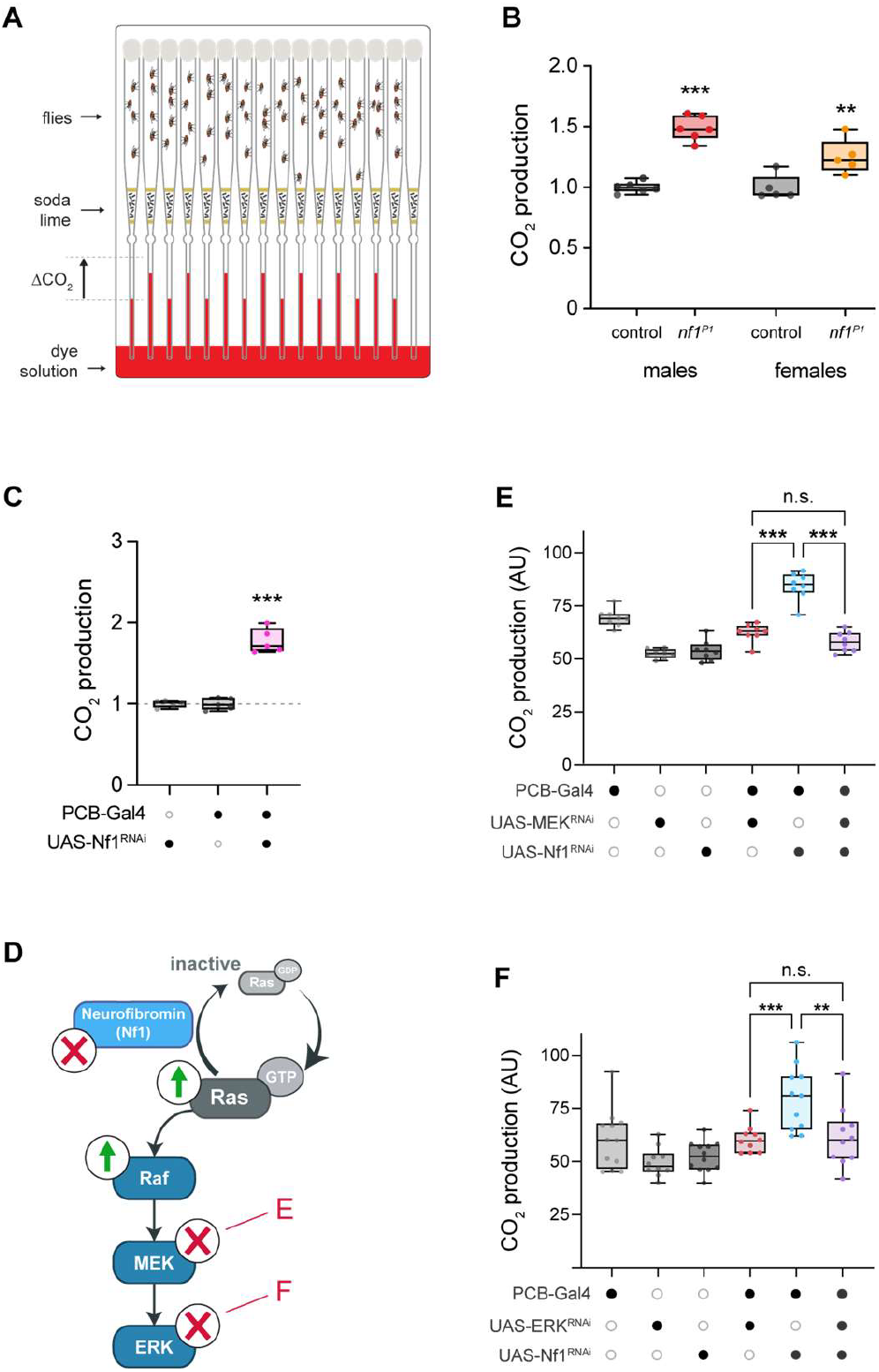
Loss of Nf1 affects metabolism via a neuronal MEK/ERK-dependent mechanism. **(A)** Diagram of the respirometry setup used to measure metabolic rate via CO_2_ production. **(B)** Normalized CO_2_ production in genomic *nf1*^*P1*^ mutant and wCS10 control males and females. **p < 0.05, ***p < 0.001 (Šidák, n = 6 males; n = 5 females). **(C)** Normalized CO_2_ production in flies with Nf1 knockdown in metabolism-regulating PCB-Gal4+ neurons compared to heterozygous Gal4/+ and UAS/+ controls. ***p < 0.001 re: both controls (Šidák, n = 5). **(D)** Diagram showing the MAPK arm of the Nf1/Ras signaling pathway. Green arrows represent the direction of signaling change following loss of Nf1. Red X marks show the molecules targeted for *in vivo* genetic analyses in panels E-F. **(E)** CO_2_ production (arbitrary units; AU) in flies harboring RNAi targeting Nf1, MEK, or both, along with controls. ***p < 0.001; n.s.: not significant (Šidák, n = 8). **(F)** CO_2_ production in flies harboring RNAi targeting Nf1, ERK, or both, along with controls. **p < 0.01, ***p < 0.001; n.s.: not significant (Šidák, n = 10-12).

The mechanistic target of rapamycin (mTOR) is a wellstudied nutrient sensor and metabolic regulator [56]. It is downstream of Nf1 and activated by Ras-PI3K-Akt (Fig. 3A). Alterations in Akt and mTOR modulate growth and cancer phenotypes in NF1 [57–60]. To test the role of mTOR, and its activators/effectors in Nf1-dependent metabolic regulation, we carried out double knockdown experiments as above. There are multiple PI3K genes (in *Drosophila*, Pi3K92E, Pi3K59F, Pi3K68D, and Pi3K21B) and some PI3K isoforms can functionally compensate for one another [47]. Therefore, we targeted the loss-of-function manipulation one step downstream, to Akt, which exists as a single isoform in *Drosophila* [48]. Knocking down Akt on its own did not affect metabolic rate (Fig. 3B). However, when Akt and Nf1 knockdown were combined, the Akt knockdown occluded the metabolic effect of Nf1 knockdown (Fig. 3B). Thus, in contrast to the behavioral phenotype, the Nf1 metabolic phenotype was dependent on Akt. Given this positive result with Akt, we moved further down the mTOR signaling pathway, testing Raptor, an mTOR complex 1 obligate subunit. Knocking down Raptor did not affect metabolism on its own, but, similar to Akt, occluded the Nf1 effect on metabolism (Fig. 3C). mTOR is a protein kinase that plays key roles in cell growth and metabolism, in part via regulating protein synthesis [61]. Activation of mTOR is positively coupled to protein synthesis via activation of S6K and inhibition of the translational repressor 4E-BP [56, 62]. Therefore, we tested whether targeting one or both of these downstream effectors could modulate the Nf1 metabolic effect. Knocking down both S6K (Fig. 3D) and 4E-BP (Fig. 3E) occluded the Nf1 metabolic effect. This demonstrated that targeting mTOR complex 1 effectors ameliorated the Nf1 metabolic phenotype.

**Figure 3:**
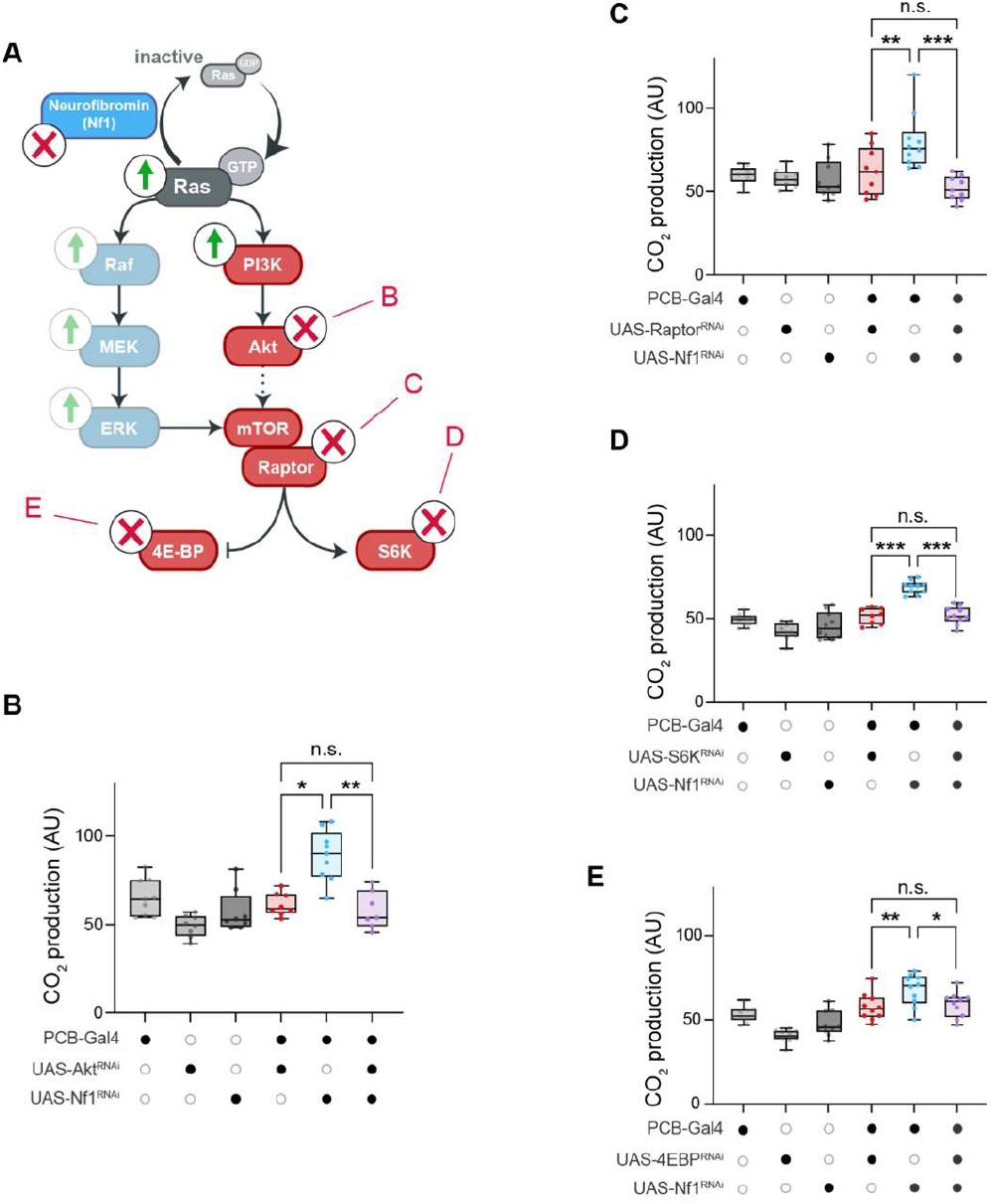
Nf1 metabolic effects require mTOR in addition to MAPK signaling mechanisms. **(A)** Diagram highlighting the mTOR arm of the Nf1/Ras signaling pathway. Green arrows represent the direction of signaling change following loss of Nf1. Red X marks show the molecules targeted for in vivo genetic analyses in panels B-E. **(B)** CO_2_ production (arbitrary units; AU) in flies bearing an RNAi targeting Nf1, Akt, or both, along with controls. *p < 0.05, **p < 0.01; n.s.: not significant (Dunn’s test, two-sided; n = 8-9). **(C)** CO_2_ production in flies bearing an RNAi targeting Nf1, Raptor, or both, along with controls. **p < 0.01, ***p < 0.001; n.s.: not significant (Šidák, n = 9-10). **(D)** CO_2_ production in flies bearing an RNAi targeting Nf1, S6K, or both, along with controls. ***p < 0.001; n.s.: not significant (Šidák, n 8-10). **(E)** CO_2_ production in flies bearing an RNAi targeting Nf1, 4E-BP, or both, along with controls. *p < 0.05, **p < 0.01; n.s.: not significant (Šidák, n = 10).

These findings reveal several key conclusions. First, Nf1 affects metabolism through coordinated actions of MAPK and mTOR signaling. Second, the Nf1 metabolic effect is susceptible to modulation of either pathway. This starkly contrasts with the behavioral phenotype, which is normalized by genetic manipulation of MEK but not Akt. Third, the metabolic effect can be rescued by downregulating any one of multiple nodes within the mTOR signaling pathway, including multiple molecules along the vertical axis of the pathway (including Akt, Raptor, and their downstream targets), as well as at least two independent parallel nodes within the pathway (S6K and 4E-BP). Thus, the metabolic phenotype is highly sensitive to any one of multiple perturbations that push signaling back toward normal levels.

### Neurofibromin mediates metabolic effects via Ras signaling independent of cAMP

Loss of Nf1 decreases cAMP and PKA levels in flies (and other model systems) (Fig. 4A) [15, 17, 20, 51, 63–65]. Therefore, we tested whether reduced cAMP levels contribute to the metabolic phenotype. As a first pass, we knocked down the type-1 adenylyl cyclase Rutabaga [66] with RNAi expressed in PCB+ neurons. This did not significantly alter metabolic rate relative to the heterozygous Gal4/+ or UAS/+ controls (Fig. 4B). This suggested that dysregulation of cAMP signaling in metabolism-controlling neurons did not mimic the Nf1 metabolic phenotype. In a complementary approach, we moved one step downstream in the signaling pathway, knocking down the PKA catalytic subunit 1 (PKA-C1). Knocking down PKA-C1 did not mimic the Nf1 metabolic effect (Fig. 4C), further suggesting that reducing cAMP/PKA signaling does not produce the metabolic effect observed in Nf1 loss-of-function conditions. A previous study reported that expression of Nf1 with a patient-derived mutation in the Ras GAP-related domain of Nf1 failed to rescue the Nf1 metabolic effects [11], initially implicating Ras signaling as the potential mechanism. The data above show that knocking down MAPK or mTOR pathways downstream of Ras can rescue the Nf1 effects on metabolism (Figs. 2, 3), and that reducing cAMP/PKA levels does not phenocopy the Nf1 effect (Fig. 4). Thus, Nf1 affects metabolism via a neuronal mechanism that depends on Ras and downstream pathways that are likely independent of cAMP/PKA.

**Figure 4:**
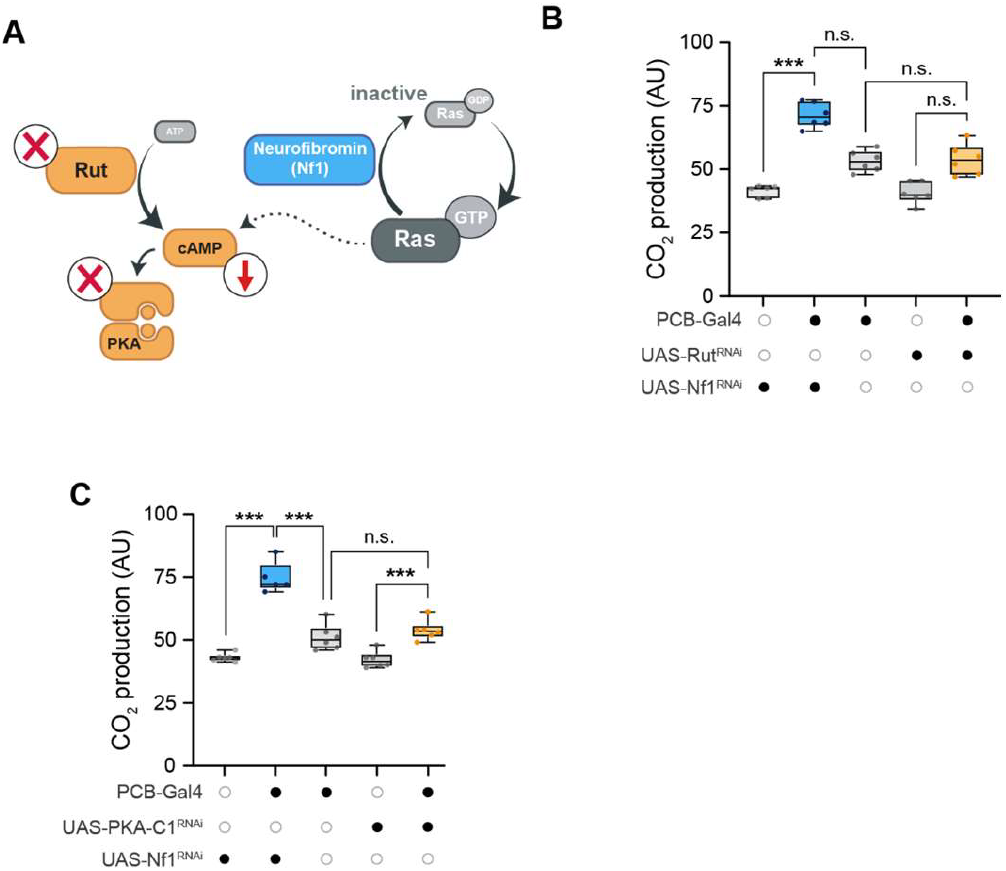
Manipulating cAMP and PKA enzyme expression did not mimic Nf1 effects on metabolic rate. **(A)** Diagram showing Nf1 and its connection to cAMP generation and PKA activity. **(B)** CO_2_ production (arbitrary units; AU) in flies expressing RNAi targeting the Rut adenylyl cyclase or Nf1 in metabolism-regulating PCB-Gal4+ neurons. ***p < 0.001; n.s.: not significant (Dunn’s test, two-sided; n = 6). **(C)** CO_2_ production in flies expressing RNAi targeting the catalytic PKA-C1 subunit or Nf1 in PCB-Gal4+ neurons. ***p < 0.001; n.s.: not significant (Šidák, n = 5-6).

### Modulation of mTOR/S6K rescues Nf1 behavioral effects independent of Akt

The above data revealed that behavioral effects of Nf1 knockdown were occluded by secondary loss of MEK but not Akt. In contrast, metabolic effects were occluded by both MAPK and mTOR pathway manipulations. There is significant crosstalk between signaling pathways, including ERK and mTOR [67] (Fig. 5A). Moreover, mTOR is a central metabolic regulator that is activated by multiple signaling pathways such as AMPK [56, 62]. We therefore asked whether the behavioral effects of loss of Nf1 could be rescued by modulating signaling at/below the level of mTOR, where multiple signaling pathways converge (Fig. 5A). To address this, we first tested whether knockdown of Raptor occluded the Nf1 effect on grooming. Raptor knockdown alone did not affect behavior but double knockdown with Nf1 occluded the effect (Fig. 5B). Next, we moved one step downstream in the pathway, testing whether S6K occluded the Nf1 effect on grooming. Double knockdown of Nf1 and S6K also occluded the effect of Nf1 knockdown alone (Fig. 5C). Thus, knocking down either Raptor or S6K rescued behavior in an Nf1 deficient background. Since MEK, but not Akt, was necessary for behavioral rescue, this suggests that either crosstalk between the pathways (Fig. 5A) could be responsible or additional inputs to mTOR (e.g., AMPK) could modulate behavioral responses independent of Akt.

**Figure 5:**
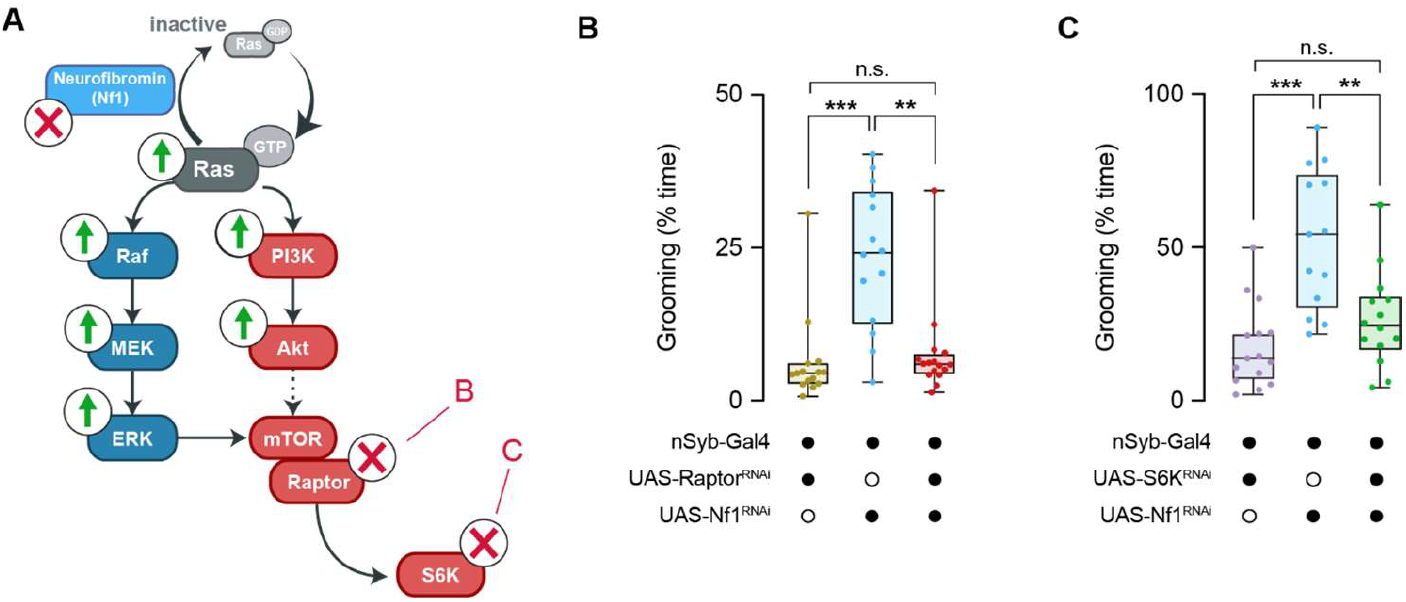
Behavioral effects of Nf1 rely on signaling downstream of mTOR. **(A)** Diagram showing Nf1-Ras signaling and the downstream MAPK and mTOR signaling pathways. Green arrows represent the direction of signaling change following loss of Nf1. Red X marks show the molecules targeted for *in vivo* genetic analyses in panels B-C. **(B)** Quantification of grooming in flies harboring RNAi targeting Nf1, Raptor, or both. **p < 0.01, ***p < 0.001; n.s.: not significant (Dunn’s test, two-sided; n = 14-16). **(C)** Quantification of grooming in flies harboring RNAi targeting Nf1, S6K, or both. **p < 0.01, ***p < 0.001; n.s.: not significant (Šidák, n = 13-15).

### Loss of Nf1 modulates metabolic rate via interneurons, with additional contributions from muscle

After establishing the downstream signaling of Nf1 for behavioral and metabolic regulation, we turned to the question of where Nf1 acts to produce effects, particularly in terms of metabolic effects. Loss of Nf1 in neurons increases CO_2_ production, and the PCB-Gal4 driver is the most restricted driver that produces a phenotype when used to knock Nf1 down [11]. Yet the driver expresses in more tissues than just neurons (the driver exhibits robust fat body expression [68], though this does not drive the phenotype [11]); whether additional tissues contribute to the Nf1-dependent change in metabolic rate has not been ruled out. To explore the potential roles of various tissues on Nf1-dependent metabolic modulation, we knocked Nf1 down in a variety of metabolically relevant tissues with RNAi. Knockdown of Nf1 in neurons with the PCB-Gal4 driver increased metabolic rate (Fig. 2C), and this is the narrowest driver identified to date [11]. The driver contains several distinct subsets of neurons, as well as additional tissues in adult animals (Fig. 6). To identify which cells are responsible for the metabolic effect, we first parsed the neuronal subsets labeled by the driver, labeling cells with mCD8::GFP (Fig. 6A-C). These include sensory neurons from the halteres and wings (Fig. 6B), interneurons in the central brain and ventral nervous cord (VNC) (Fig. 6A), and insulin-producing cells [11]. A prior study found that loss of Nf1 in insulin-producing cells does not affect metabolism [11], leaving sensory neurons and interneurons as remaining candidates. To test the role of sensory neurons, we knocked down Nf1 in neurons innervating the campaniform sensillae (CS), a type of sensory neurons in the wings and halteres (Fig. 6B). The DB331-Gal4 driver labels haltere CS afferents [69] and knocking down Nf1 with this driver did not detectably alter metabolic rate (Fig. 6E). In a complementary experiment, we knocked Nf1 down in wing campaniform sensillae using R12C07-Gal4 [69]. Similar to DB331, we did not detect a difference in metabolic rate when knocking Nf1 down with R12C07 (Fig. 6E). This suggests that while the PCB-Gal4 driver drives expression in campaniform afferents, these neurons are not responsible for the alterations in metabolic rate. This leaves interneurons in the central nervous system as the likely cell-autonomous neuronal source of the neuronal Nf1 metabolic effect.

**Figure 6:**
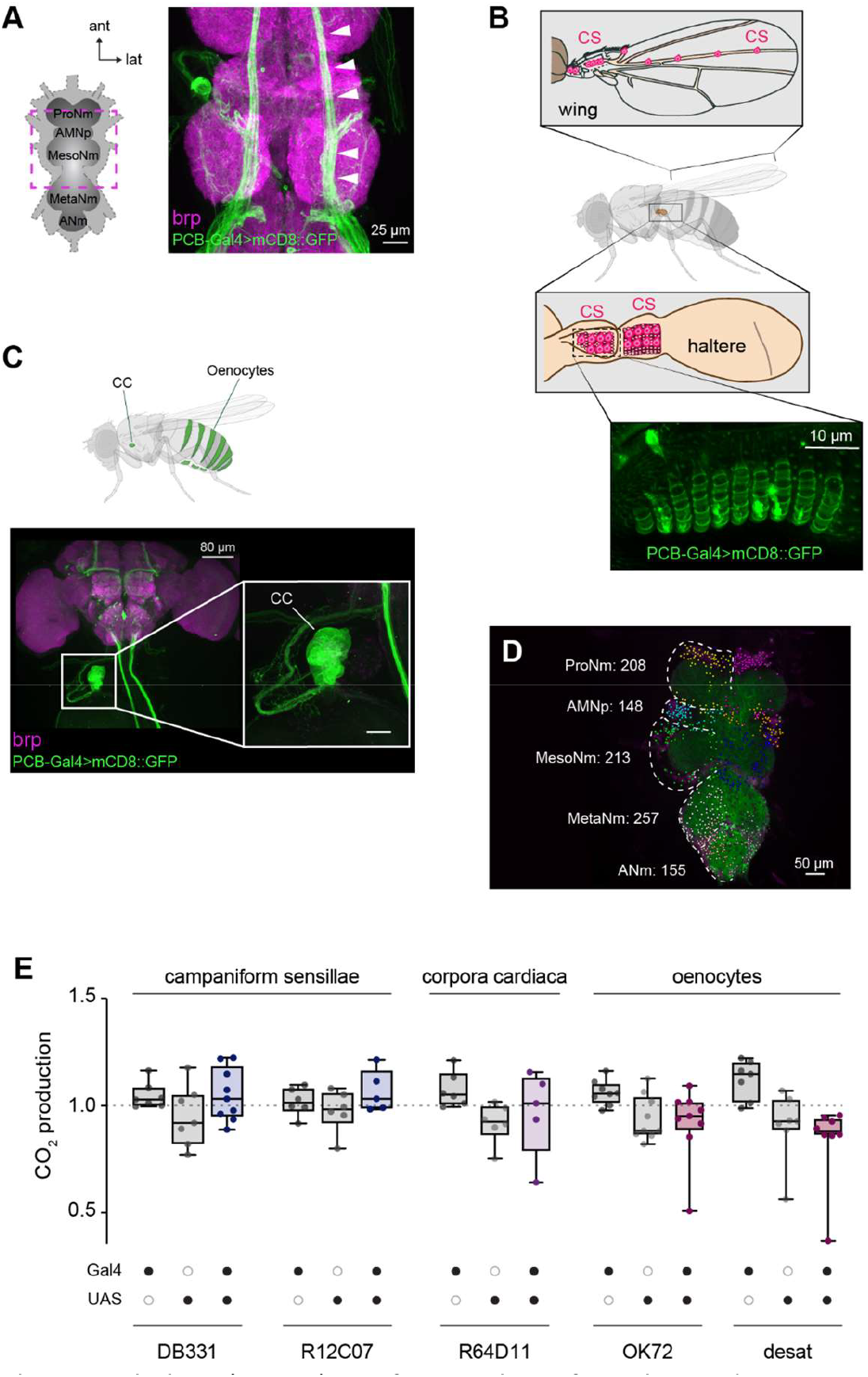
Neurons and other tissues labeled by the PCB-Gal4 driver that potentially contribute to Nf1 metabolic effects. **(A)** Diagram and GFP image of the haltere tract passing through the ventral nerve cord (VNC). Left: diagram of the VNC. ProNm: prothoracic neuromeres, AMNp: accessory metathoracic neuromeres, MesoNm: mesothoracic neuromeres, MetaNm: metathoracic neuropil, ANm: abdominal neuromere. Ant: anterior, lat: lateral. Right: immunostained VNC from a PCB-Gal4>UAS-mCD8::GFP fly counterstained with the neuronal marker bruchpilot (brp). **(B)** Diagram of campaniform sensillae (CS) in the wing and haltere, along with a GFP image showing that PCB-Gal4 labels CS in the haltere. **(C)** Diagram showing the location of the corpora cardiaca (CC) and oenocytes, along with an image of the PCB-Gal4 driver showing labeling in the brain and adjacent corpora cardiaca (CC). The tissue was counterstained with brp. Scale bars = 80 μm. **(D)** VNC with PCB-Gal4+ cells labeled with nuclear-localized GFP.nls. Cell bodies were counted and marked with dots. Each neuromere is outlined with dashed white lines the left hemisphere, and the number of labeled cells is noted on the left side. **(E)** Quantification of CO_2_ production when Nf1 was knocked down using drivers expressing in CS sensory neurons (DB331-Gal4 [p = 0.19, ANOVA, n = 7-9], R12C07-Gal4 [p = 0.26, ANOVA), CC (R64D11-Gal4 [p = 0.14, ANOVA, n = 5-6]), or oenocytes (OK72-Gal4 [p = 0.03, Kruskal-Wallis, n = 8-9], desat1-Gal4 [P < 0.001, Kruskal-Wallis, n = 7-8]).

Next, we examined PCB-Gal4 for any non-neuronal Nf1 metabolic contributions. In addition to the brain and ventral nervous system, the Gal4 driver strongly labels neurosecretory cells in the corpora cardiaca (CC) (Fig. 6C). A previous study reported that knocking Nf1 down with either of two CC drivers, Akh-Gal4 and Tk-Gal4 did not affect metabolic rate [11]. Nonetheless, given the strong expression of PCB-Gal4 in the CC, we further challenged the role of this driver in Nf1-dependent metabolic regulation here. Knocking down Nf1 with an additional CC-selective Gal4 driver, R64D11-Gal4, did not alter metabolism (Fig. 6E), confirming that Nf1 in the CC does not cell-autonomously regulate metabolism. Beyond the CC, additional cell types regulate metabolism, including the fat body and oenocytes (Fig. 6C). The fat body is a primary lipid storage site, with similar function to mammalian adipose tissue [70], yet Nf1 knockdown in the fat body does not affect metabolism [11, 68]. Oenocytes are hepatocyte-like neurosecretory cells in the fly abdomen that respond to nutrient state, synthesize very long-chain fatty acids, and regulate energy homeostasis [71] (Fig. 6C). Knocking Nf1 down with either of two oenocyte-expressing drivers, OK72 and desat, did not significantly affect metabolic rate (Fig. 6E). There was a slight decrease in the experimental group relative to controls, resulting in a main effect in the ANOVA. However, the experimental group did not significantly differ from both Gal4/+ and UAS/+ groups using either driver. Furthermore, the trend was in the downward direction and therefore could not account for the Nf1 effect on metabolism. Thus, we conclude that Nf1 does not impact metabolic rate via functions in the CC, fat body, or oenocytes.

The above data implicate neurons in the PCB-Gal4 driver. This driver labels a set of neurons in the central brain and VNC [11]; the metabolic phenotype is suppressed by the ventral nervous system-biased tsh-Gal80 repressor [11], suggesting that the VNC is a likely locus of Nf1 metabolic effects. To gain insight into the neurons that could be responsible for the phenotype, we labeled nuclei of PCB-Gal4+ cells with nuclear-localized GFP (GFP.nls) and counted the labeled cells in the VNC (Fig. 6D). There were 981 labeled cell bodies across the VNC, with 208 in the prothoracic neuromeres (ProNm), 148 in the accessory metathoracic neuromeres (AMNp), 213 in the mesothoracic neuromeres (MesoNm), 257 in the metathoracic neuropil (MetaNm), and 155 in the abdominal neuromere (ANm) (Fig. 6D). We observed no evidence of glial-like labeling in mCD8::GFP labeled brain (Fig. 6C) or VNC (Fig. 6A), so these GFP.nls labeled cell bodies likely represent neuronal somata. The VNC contains approximately 23,000 neurons [72], so the 981 labeled neurons in the PCB-Gal4 driver represent approximately 4.3% of the total neurons that have cell bodies within the VNC. Overall, this suggests that PCB-Gal4+ non-neuronal cells (oenocytes and corpora cardiaca), neurons innervating the VNC (campaniform sensillae afferents in the haltere tract), as well as other candidates (insulin-producing cells, peptidergic cells, and fat body [11]) are not responsible for the Nf1 effect on metabolism. Thus, interneurons within the VNC are the likely major modulators of Nf1-dependent neuronal metabolic effects.

Muscle tissue is an additional candidate for non-neuronal Nf1 effects on metabolism [30, 35, 73]. Muscle movement and maintenance are energetically costly – in insects, skeletal (flight) muscles consume the largest amount of energy in proportion to the animal’s weight [74, 75]. Additionally, Nf1 exerts effects on metabolic parameters in mammalian muscle tissue [30, 38, 73, 76]. To test whether loss of Nf1 in muscle contributes to the organismal metabolic phenotype, we knocked Nf1 down using four Gal4 drivers that include muscle expression: Mef2, c179, 24B, and R22H05. Metabolic rate was increased when Nf1 was knocked down with Mef2, c179, and R22H05, but not 24B. Among these drivers, 24B is expressed in muscle broadly through development, but its expression drops progressively after eclosion [77]. Therefore, the lack of effect when knocking Nf1 down with 24B suggests that Nf1 could impact metabolism via post-developmental effects in adult flies, though differences in the expression patterns between the drivers (across muscle or off-target outside muscle) could also be responsible. The effect size of the metabolic phenotype when knocking Nf1 in muscle (Fig. 7I) was smaller than when knocking down Nf1 with the PCB-Gal4 driver (Fig. 2C) or pan-neuronal knockdown [11]. This suggested that the muscle contributions are either secondary to neuronal contributions or due to off-target expression of Gal4 in neurons within the muscle-selective Gal4 drivers. To test whether the effect of Nf1 knockdown using these drivers was due solely to muscle or a combination of muscle and neurons, we used a muscle-selective MHC-Gal80 to repress Mef2-Gal4 expression in muscle. This manipulation only marginally suppressed the Mef2-Gal4 phenotype – metabolic rate in flies expressing Nf1RNAi under control of the MHC-Gal80;Mef2-Gal4 combination were not significantly different from either the Mef2-Gal4>Nf1RNAi or the heterozygous controls (Fig. 7J). Thus, the metabolic effect may be dominated by neuronal expression in the driver(s); for instance, Mef2-Gal4 contains robust expression in the brain (along with skeletal and visceral muscle) [78]. Overall, these data suggest that interneurons within the VNC are the major source of the Nf1 metabolic effect, with additional contribution from Nf1 in muscle.

**Figure 7:**
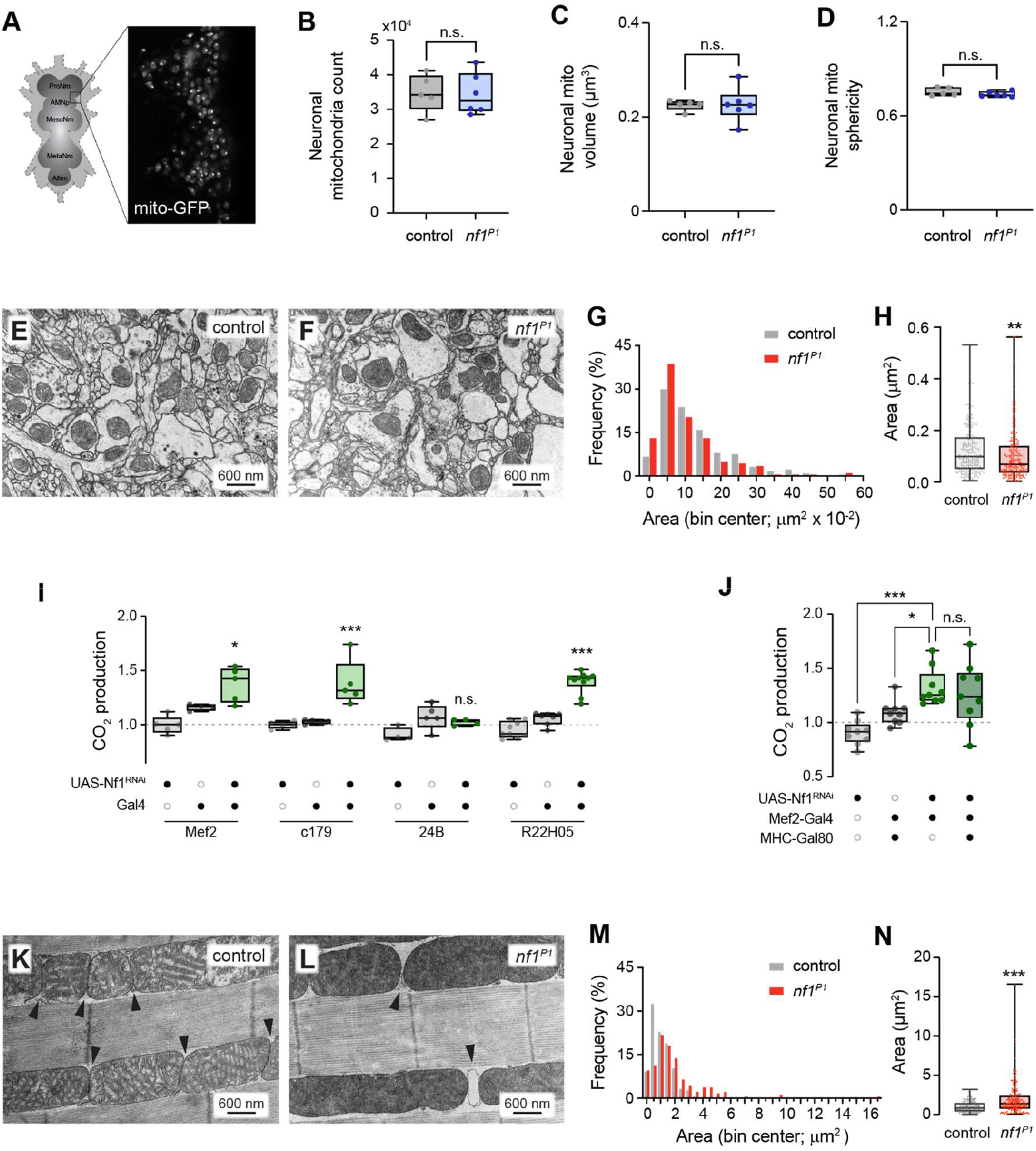
Nf1 effects in muscle contribute to the metabolic alterations. **(A)** Diagram of ventral nerve cord with neuronal mitochondria labeled with GFP. **(B)** Neuronal mitochondrial counts in control flies and *nf1*^*P1*^ mutants (unpaired t-test; n = 5-6). **(C)** Neuronal mitochondrial volume in control flies and *nf1*^*P1*^ mutants (unpaired t-test; n = 5-6). **(D)** Neuronal mitochondrial sphericity in control flies and *nf1*^*P1*^ mutants (unpaired t-test; n = 5-6). **(E)** Representative transmission electron micrograph of protocerebral neuropil from a control (wCS10) fly. **(F)** Representative transmission electron micrograph of protocerebral neuropil from an *nf1*^*P1*^ mutant fly. **(G)** Histogram of neuronal mitochondria size, quantified from electron micrographs from the *Drosophila* protocerebrum. Measurements are 2D cross-sectional area in μm^2^, graphed in 0.05 μm^2^ width bins (n = 227 [wCS10], 207 [*nf1*^*P1*^]). **(H)** Individual mitochondria area measurements, quantified from electron micrographs (same values plotted in panel G). **p < 0.01 (Mann-Whitney). **(I)** Knockdown of Nf1 with certain muscle-selective Gal4 drivers. *p < 0.05, ***p < 0.001 (Šidák, n = 4-10). **(J)** Knockdown of Nf1 with the muscle selective Gal4 driver Mef2-Gal4, with and without suppression of Gal4 expression in muscle via the MHC-Gal80 (Šidák, n = 9). **(K)** Representative transmission electron micrograph of flight muscle from a control (wCS10) fly. Junctions between adjacent mitochondria are highlighted with black arrowheads. **(L)** Representative transmission electron micrograph of flight muscle from an *nf1*^*P1*^ mutant fly. Spacing between adjacent mitochondria is highlighted with black arrowheads. **(M)** Histogram of muscle mitochondria size, quantified from electron micrographs from the *Drosophila* flight muscle. Measurements are 2D cross-sectional area in μm^2^, graphed in 0.5 μm^2^ width bins (n = 185 [wCS10] and 189 *nf1*^*P1*^]). **(N)** Individual mitochondria area measurements, quantified from electron micrographs (same values plotted in panel M). ***p < 0.001 (Mann-Whitney).

### Nf1 deficiency did not detectably alter neuronal structure or mitochondria

Alterations in metabolism could result from changes in mitochondrial structure and/or function. Loss of Nf1 impacts ROS production [41] and mitochondrial bioenergetics in an oncogenic context [31]. To test whether mitochondria were altered, we first labeled neuronal mitochondria pan-neuronally with a genetically-encoded, mitochondrial-targeted fluorescent protein, and analyzed mitochondria in the VNC (Fig. 7A-D). At the light microscopy level, we detected no significant differences in mitochondria number between controls and *nf1*^*P1*^ mutants (Fig. 7B). There were also no differences in mitochondrial volume (Fig. 7C) or sphericity (Fig. 7D) between *nf1*^*P1*^ mutants and controls. Analysis of electron micrographs taken from the protocerebrum neuropil area were unremarkable, revealing no major qualitative differences between *nf1*^*P1*^ mutants and wCS10 controls with respect to overall neuropil morphology, synaptic T-bar and vesicle distribution and size, and membrane integrity (Fig. 7 E,F). Quantification of mitochondrial area revealed a decrease in mitochondria size in protocerebral neurons (Fig. 7 G,H). While this difference was statistically significant, the magnitude was modest, and there was little shift in the distribution of mitochondria sizes (Fig. 7G). Thus, these data suggest that neither mitochondrial regulation (e.g., via fission/fusion) nor neuronal ultrastructure were detectably altered in *nf1* mutant neurons.

### Altered muscle mitochondrial morphology in *nf1* mutants

As loss of Nf1 affected metabolism via effects in muscle (as well as neurons) (Fig. 7I), we asked whether muscle morphology and mitochondria were affected by Nf1 deficiency. To test this, we examined electron micrographs of *nf1*^*P1*^ mutants and controls. Ultrastructural analysis of *Drosophila* flight muscle revealed qualitative and quantitative changes in mitochondria. There was visibly abnormal spacing between adjacent mitochondria in *nf1*^*P1*^ mutant muscle (Fig. 7 K,L). Further, the mitochondria were larger in *nf1*^*P1*^ mutants, exhibiting an increase in cross-sectional area relative to wCS10 controls, which was accompanied by a rightward shift in the distribution in the mutants (Fig. 7 M,N). Thus, there were robust changes in muscle mitochondria ultrastructure. Overall, loss of Nf1 affected neurons and muscle differently at the ultrastructural level, with muscle exhibiting notable alterations in mitochondrial structure.

## Discussion

Neurofibromatosis type 1 is a monogenetic disorder that affects diverse and complex behavioral, cellular, and organismal phenotypes. While the disorder results from mutations in a single gene, its protein product regulates a variety of diverging downstream signaling pathways. This raises the question of whether multiple neurofibromatosis type 1 phenotypes exhibit similar or differing dependence on the signaling pathways downstream of Nf1 and Ras. To test this, we implemented *in vivo* genetic analysis of signaling pathways in *Drosophila* - a robust platform for dissecting the action of individual signaling molecules on organismal phenotypes. Two human disorder-relevant organismal phenotypes – behavior (grooming) and metabolic modulation – exhibited differing dependence on signaling pathways immediately downstream of Nf1/Ras. Nf1-dependent behavioral alteration was rescued by manipulation of MEK but not Akt. In contrast, the Nf1 metabolic phenotype was rescued by both MEK and Akt (Fig. 8).

**Figure 8:**
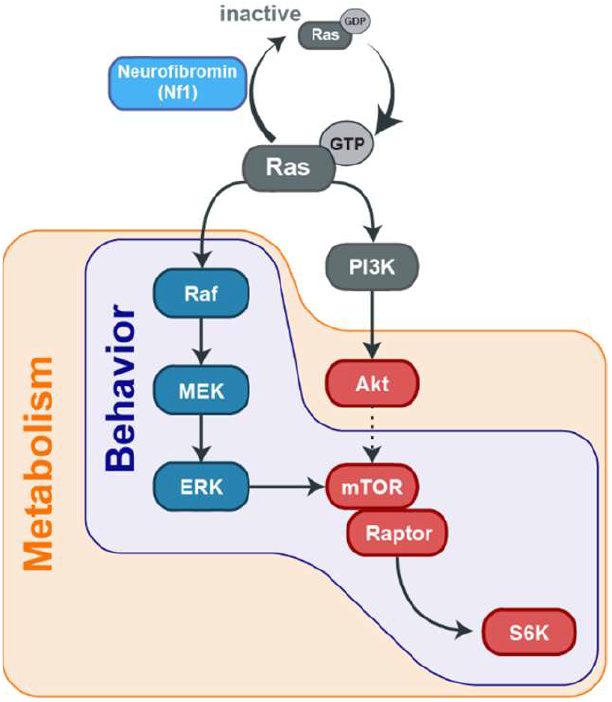
Summary diagram showing Nf1-Ras signaling and the downstream MAPK and mTOR signaling pathways. The signaling pathways implicated in behavioral and metabolic signaling are highlighted. Molecules that were tested with double RNAi in both assays are shown (Pi3K is included as an upstream signaling molecule in the mTOR pathway but was not tested).

The Nf1 metabolic phenotype exhibited striking reliance on multiple signaling molecules. These were spread across at least two parallel signaling pathways (MAPK and mTOR), included hits distributed vertically along the same pathways (MEK/ERK, Akt/Raptor), and included one pair of molecules situated in parallel within the same pathway (S6K/4E-BP). Despite the breadth of these hits across multiple pathways/loci, each molecule was individually necessary for the Nf1 effect on metabolism – knocking any of them down occluded the Nf1 effect on metabolism. This suggests that Nf1-dependent metabolic modulation results from coordinated actions of MAPK (Raf-MEK-ERK) and mTOR (PI3K-Akt-mTOR-S6K/4E-BP) signaling. Considering the pathways examined in the present study, the Nf1 metabolic phenotype appears to be dependent on a wider range of signaling molecules than the behavioral (grooming) phenotype.

A major difference in pathway dependency between the behavioral and metabolic phenotypes was Akt - behavioral effects of Nf1 knockdown were normalized by knocking down MEK but not Akt (Fig. 8). This suggests that Nf1 exerts behavioral effects via more selective effects on MAPK signaling at high levels of the signaling cascade (proximal to Ras). Within the PI3K/Akt/mTOR signaling pathway, there are multiple isoforms of signaling molecules at several levels of the pathway. For instance, in *Drosophila*, there are multiple PI3K proteins/isoforms (PI3K92E/Dp110, PI3K68D/cpk, PI3K59F/Vps34, and Pi3K21B), like mammals. This complicates analysis with loss-of-function approaches, as different isoforms could compensate one another. Therefore, we focused our analysis on Akt and mTOR. In mammals, there are three isoforms of Akt (AKT1, AKT2, and AKT3), and knockout of each can produce differing effects [79]. In *Drosophila*, there is only one Akt gene, facilitating unambiguous genetic dissection of the role of Akt in cellular signaling. Here we leveraged this to test the role of Akt in mediating Nf1 effects.

Targeting deep levels of the mTOR signaling pathway (Raptor and S6K) normalized both Nf1 behavioral effects and metabolism. Thus, Akt was not required for the Nf1 behavioral effect, but mTOR and S6K were. This suggests that mTOR complex 1 and S6K contribute to the behavioral phenotype but are pathologically hyperactivated by Nf1 in an Akt-independent manner. Crosstalk between MAPK and mTOR signaling pathways is one way in which this could occur. Multiple points of crosstalk between these pathways have been reported, including ERK-mediated phosphorylation of Raptor and cross-activation via intermediates such as RSK and TSC1/2 (Fig. 8) [67]. The existence of such crosstalk facilitates potential treatment of NF1 behavioral phenotypes in several ways. For instance, it could be possible to target a small set of upstream signaling molecules (e.g., MEK) or any one of a larger set of downstream signaling molecules representing their direct targets (e.g., ERK) or indirect targets (e.g., mTOR). Further preclinical studies will be needed to identify viable approaches.

Nf1’s metabolic effects were Akt-dependent, which likely functioned through activation of downstream mTOR signaling. Akt activates the Raptor-containing mTOR complex 1 via a signaling pathway involving TSC1/2 and Rheb [61]. mTOR complex 1 is a conserved central metabolic regulator [80], exerting effects on cellular growth and metabolic processes [61]. The dependence of the Nf1 metabolic phenotype on both MAPK and Akt/mTOR signaling pathways is reminiscent of signaling in various cancers. For instance, both MEK and Akt are hyperactivated in optic gliomas, and inhibition of either pathway reduces several optic glioma phenotypes [59]. Similarly, inhibition of Akt and MEK synergistically suppresses NF1 malignant peripheral nerve sheath tumor (MPNST) growth [81]. The cellular mechanism remains to be fully elucidated, though it could involve metabolic alterations; Akt promotes mitochondrial fusion and elongation [82]. Whether this represents a similar mechanism to that driving alterations in organismal metabolic alterations is unclear – we detected no major changes in neuronal morphology, though there was evidence for changes in muscle mitochondria.

Multiple signaling pathways have been implicated in neurofibromatosis type 1 phenotypes, including Ras/MAPK, Ras/PI3K/mTOR, and cAMP/PKA. The commonly-studied MAPK signaling pathway is implicated in multiple phenotypes, and MEK is the target for current FDA-approved treatments [21, 22]. Nf1 also regulates cAMP in a Ras-dependent manner via the activation of atypical protein kinase C (PKCζ) in neurons [83]. In the present study, we did not find evidence for cAMP effects on the Nf1 metabolic effect, as cAMP/PKA manipulations did not phenocopy the nf1 mutant phenotype. One caveat is that – given this lack of phenocopy – we did not attempt to normalize cAMP/PKA levels in the mutant background. mTOR is hyperactivated in neurofibromatosis type 1 and regulates some neurofibromatosis type 1 phenotypes, particularly tumors [84, 85]. Both Ras/ERK and cAMP/PKA signaling regulate aspects of cognition in animal models [63, 86, 87]. Ras signals through a variety of additional effectors, including RalGEF/RalA, TIAM1, PLC, RASSF1/5. Several of these pathways have been implicated in neurofibromatosis type 1 tumor phenotypes. RalGEF/RalA is a Ras effector pathway that is overactivated in NF1 malignant peripheral nerve sheath tumors (MPNST) [88, 89]. RASSF1 expression is altered in NF1 MPNST via DNA promoter methylation-induced silencing, and this is associated with poor prognosis [90]. Ras-TIAM1-Rac signaling is implicated in proliferation of plexiform neurofibromas, as genetic disruption of Rac prevents tumor formation [91]. Combination therapies targeting multiple pathways may be an effective strategy both for cancers [92, 93] and other phenotypes, such as the cognitive and behavioral symptoms.

At the circuit/systemic level, Nf1 exerts a primary effect in neurons to modulate the Nf1 metabolic phenotype, with a secondary contribution from muscle tissue. The critical neurons are some set of the 981 VNC interneurons labeled by the PCB-Gal4 driver. In addition to the interneurons the PCB-Gal4 driver labels multiple other cell types, including campaniform sensillae in the halteres and wings, corpora cardiaca, and fat body. Tissue-specific RNAi experiments in the present and a prior study [11] revealed no effect of knocking Nf1 down in these cell types, suggesting that these tissues may not contribute. Oenocytes represent an additional potential site of systemic metabolic effects, which we tested here. Caveats to these experiments include the possibility of potential shifts in Gal4 expression patterns of the drivers over development/lifespan [94], developmental effects of Nf1 on neurons [43], “off-target” expression sites [95], or nonlinear expression levels across labeled cells. To minimize the potential for these confounds, we have tested multiple lines covering each of these cells/tissues. Thus, multiple lines of evidence support a role for neurons and muscle in mediating Nf1 effects on metabolism.

In addition to the neuronal effects of Nf1, knocking down Nf1 with muscle-selective Gal4 drivers altered metabolism and increased CO_2_ production. Suppressing muscle expression of the Mef2-Gal4 driver with MHC-Gal80 did not significantly affect the metabolic rate, reinforcing the interpretation that muscle is not the primary/only source of the Nf1 metabolic effect. Subcellular changes in nf1 mutants included differences in mitochondrial ultrastructure between *nf1* mutants and wild-type controls. In the nervous system, there were no major morphological differences in neurons between nf1 mutants and controls in terms of overall neuronal morphology or synaptic structure, though there was a small shift in the distribution of mitochondrial size. In muscle, there was a more pronounced shift in mitochondrial size, which was accompanied by a qualitative change in mitochondrial appearance (increased spacing). This change in mitochondrial appearance was subtle, yet readily detectable in *Drosophila* flight muscle since the mitochondria line up end-to-end. In comparison, mitochondria in mice with biallelic Nf1 inactivation in muscle appear normal, though there is significant muscle atrophy, metabolic dysregulation, and ultimately lethality [30]. The mitochondrial changes in *Drosophila* were reminiscent of swollen mitochondria reported in another study [96]. Loss of nf1 drives mitochondrial dysregulation [41], which could contribute to changes in mitochondrial regulation and morphology.

Overall, this study revealed how loss of Nf1 modulates two organismal phenotypes through effects on two major Ras effector pathways: MAPK and mTOR. The results suggest that multiple pathway nodes could be targeted to approach NF1 phenotypes, particularly the metabolic alterations. Yet some of the phenotypes exhibit differential dependence on certain signaling nodes, a feature that could be exploited to target different NF1 phenotypes independently. While MAPK and mTOR are two of the major, well-studied Ras effectors, Ras signals through a wide range of effectors. Future studies will be required to address these signaling pathways more broadly. The diversity and complexity of Ras signaling represents both a challenge and an opportunity for identifying new therapeutic targets for NF1 and other Rasopathies.

## Acknowledgements

We thank Lita Duraine and the Baylor College of Medicine Jan Duncan Neurological Research Institute microscopy facility for assistance with transmission electron microscopy. In addition, we are grateful to David Gutmann, Corina Anastasaki, Jonathan Payne, J. Elliot Robinson, Robert Kesterson, and the members of the CABIN task force for feedback on Nf1 aspects of the project, as well as Nathan Mohar and Lori Wallrath for helpful discussions on muscle components of the project. We thank Timothy Mosca for MHC-Gal80, Ronald L. Davis for UAS-4mtGCaMP3 and UAS-RFP, Michael Dickinson for DB331-Gal4, and James Walker and Andre Bernards for *nf1* mutants. This research was supported by NIH/NINDS R01 NS114403, R01 NS124716, R01 NS126361, R01 NS097237, R21 NS124198, DOD/CDMRP NF230039, and the Roy J. Carver Charitable Trust. The Leica SP8 confocal with STED capability in the University of Iowa Central Microscopy Resource Facility was supported by a grant from the Roy J. Carver Charitable Trust, with additional funding from the Office of the Vice President for Research, the Carver College of Medicine, and the College of Liberal Arts and Sciences. Stocks used in this study were obtained from the Bloomington *Drosophila* Stock Center (NIH P40OD018537). We thank Linda Buckner and Robert Svetly for administrative assistance. This preprint was typeset with the bioRxiv word template by @Chrelli: www.github.com/chrelli/bioRxiv-word-template

## Materials and Methods

### Drosophila maintenance

Flies were raised on *Drosophila* cornmeal/agar food medium according to standard protocols and housed at 25 ºC, 60% relative humidity, on a 12:12 h light:dark cycle. The *nf1*^*P1*^ mutation was backcrossed six generations into the control genetic background wCS10. RNAi lines were obtained from the Vienna *Drosophila* RNAi Center, Gal4/+ control crosses consisted of an empty attP control line (VDRC #60100). UAS-dicer2 was included to potentiate the RNAi effect [97], and was included all genotypes containing the UAS-RNAi. Male flies were used for all experiments, unless otherwise specified, to prevent egg accumulation in behavioral and respirometry chambers.

### Behavioral analysis

Spontaneous grooming was quantified in an open field [44, 98]. Individual animals were aspirated into an open field area, 15.4 mm in diameter and 2.85 mm in height, consisting of an opaque white PLA boundary wall covered on the top and bottom with clear polycarbonate sheets. The arena was illuminated from below with light from white light-emitting diodes that was diffused through a sheet of white acrylic. Light intensity was measured at 720 lm/m^2^ in the location of the fly. Videos were recorded at 7.5 frames per second, 1,616 x 1,240, lossless Motion JPEG 2000 compression using a camera mounted above the chamber (FLIR Teledyne Blackfly S) fitted with a 25 mm lens (Edmund Optics). Five-min videos were recorded 30 min following the introduction into the open-field arena. Grooming was manually scored via frame-by-frame analysis, recording the start and stop frame for each grooming bout. Grooming was calculated as the percent of total time grooming during the 5-minute video.

### Metabolic analysis

CO2 production was quantified via respirometry [11, 55]. Respirometers were constructed by gluing together a 1 ml pipette tip and 50 µl capillary micropipette. Soda lime was placed into each pipette tip between two foam pieces. Flies were sorted under CO_2_ anesthesia and allowed to recover for at least 24 h before beginning the experiment. Four flies of the same genotype were aspirated into each respirometer, and the top was sealed with non-hardening modeling clay. Pipette seal was monitored and any that leaked were excluded from analysis. Sixteen respirometers were hung on a custom-made rack in a latch-lid chamber. One control respirometer was left empty in each experiment. The bottom of the chamber was filled with a red water-based dye and the chamber lid was closed/sealed with vacuum grease. The chamber was placed in an incubator and allowed to equilibrate at 25 ⍰C for 1 h. Images were captured every 15 min with PhenoCapture 3.3. The liquid meniscus level was measured in each respirometer after 3 h using Fiji 2.0. Where appropriate, data were normalized to the mean of the control genotypes.

### Quantitative polymerase chain reaction (qPCR)

Heads were collected from adult flies (5–10 days post-eclosion; 30 heads per genotype). Each sample was added to QIAzol Lysis Reagent (Qiagen, catalog #79306) and homogenized using a mortar and pestle. RNA was isolated with the Qiagen RNeasy Lipid Tissue Mini Kit. Complimentary DNA library was constructed using LunaScript RT Master Mix (New England Biolabs, catalog #E3025L) with the Random Primer Mix (New England Biolabs, catalog #S1330S). Nf1 gene expression was quantified in fly head samples using Luna Universal qPCR Master Mix (New England Biolabs, catalog #M3003L). Primer sequences were as follows: 5’-CTTTTGGCACGTTTCGAGGAT-3’ (Nf1_F), 5’-GGTAGCGCGATATGTGGATCA-3’ (Nf1_R), 5’-ATGCTAAGCTGTCGCACAAATG-3’ (Rpl32_F), and 5’-GTTCGATCCGTAACCGATGT-3’ (Rpl32_R). Five biological replicates, with three technical replicates each, were analyzed for each genotype. Analysis of gene expression was performed using standard ΔΔCt analysis. Gene expression was normalized to Rpl32, and log2 fold change was calculated.

### Western blot analysis

For analysis of pERK/ERK/β-tubulin, lysates of five-day old adult fly heads were prepared using RIPA buffer supplemented with Pierce Protease and Phosphatase inhibitors (Thermo Scientific). Protein samples were mixed with NuPAGE 4x LDS Sample Buffer (Invitrogen), resolved on 4–15% Tris-Glycine Mini-PRO-TEAN TGX Stain-Free Protein Gels (Bio-Rad) with Tris Glycine SDS. Gels were transferred to polyvinylidene difluoride (PVDF) membranes and 5% BSA in TBST was used for blocking. Primary antibodies used were rabbit pERK (CST, 9101, 1:1000), rabbit ERK (CST, 4695, 1:1000), and mouse anti-β-tubulin (DHSB, E7, 1:10000). The membranes were washed with TBST, followed by secondary antibody incubation for 1 hour at room temperature. Secondary antibodies used: goat anti-Mouse IgG (H+L) Alexa 800 (Invitrogen, A32730, 1:10000) and donkey anti-Rabbit IgG (H+L), Alexa 680 (Invitrogen, A10043, 1:5000). Membranes were stripped using NewBlot PVDF stripping buffer (LI-COR, 928-40032) after blotting with pERK/β-tubulin, then reblocked in 5% BSA and incubated with ERK/β-tubulin antibodies overnight. Images were captured with an Odyssey XF Imaging System (LI-COR).

### Immunohistochemistry and imaging

Five to ten-day-old adult flies were dissected fixed in 1% paraformaldehyde in in Schneider’s *Drosophila* medium and processed as previously described [99]. Samples were stained with primary antibodies for 3 hours at room temperature and 4 ⍰C overnight, followed by secondary antibodies for 3 hours at room temperature and 4 days at 4 ⍰C. Incubations were performed in blocking serum (3% normal goat serum). Samples were mounted in Vectashield (Vector Laboratories) for imaging. The following antibodies were used: rabbit anti-GFP (1:1000, Invitrogen), mouse anti-brp (nc82) (1:50, DSHB), goat anti-rabbit IgG, and goat anti-mouse IgG (1:800, Alexa 488 or Alexa 633, respectively, Invitrogen). Mitochondria were fluorescently labeled by expressing UAS-4mtGCaMP3 and immunostaining the GFP subunit of the GCaMP molecule. Cellular nuclei were labeled with DAPI (1:1000, Invitrogen). A Leica SP8 confocal microscope with LAS X software was used to obtain images following standard protocols.

### Transmission Electron Microscopy and ultrastructure analysis

*Drosophila* head and flight muscle ultrastructure was imaged following standard Electron Microscopy procedures using a Ted Pella Bio Wave processing microwave with vacuum attachments. The tissue was covered in 2% paraformaldehyde, 2.5% Glutaraldehyde, in 0.1 M Sodium Cacodylate buffer at pH 7.2. After dissection the heads were incubated for 3 days in fixative in a cold room rotator. All processing from fixative to Ethanol were done on ice. The pre-fixed heads were then fixed again in the microwave vacuum processor, followed by 3x millipore water rinses, post-fixed with 1% aqueous osmium tetroxide, and rinsed again 3x with millipore water. Concentrations from 30-100% of Ethanol were used for the initial dehydration series, followed with Propylene Oxide as the final dehydrant. Samples were gradually infiltrated with 3 ratios of propylene oxide and Embed 812, finally going into 3 changes of pure resin under vacuum. Samples were allowed to infiltrate in pure resin overnight on a rotator. The samples were embedded into flat silicone molds and cured in the oven at 62 ⍰C for three days. The polymerized samples were thin-sectioned at 48-50 nm and stained with 1% uranyl acetate for ten minutes followed by lead citrate for one minute before TEM examination. Grids were viewed in a JEOL 1400 Plus transmission electron microscope at 80kV. Images were captured using an AMT XR-16 mid-mount 16 mega-pixel digital camera.

### Statistical analysis

Normality of data was assessed with the D’ Agostino Pearson Test. In figures, box plots graph the median as a line, the interquartile range (IQR) as a box, and whiskers extend to the min/max values. Hypothesis testing was carried out using t tests or ANOVA followed by Šidák’s multiple comparisons tests (parametric), or Wilcoxon rank-sum test or Kruskal-Wallis omnibus test followed by Dunn multiple comparisons tests (nonparametric). Two-way comparisons were carried out with a two-way ANOVA followed by Šidák’s multiple comparisons tests. For RNAi and Gal4-mediated over-expression, the experimental group was compared to heterozygous Gal4/+ and UAS/+ controls and considered positive only if it significantly differed from both controls in the same direction. For double RNAi experiments, the double RNAi group was compared to both single RNAi groups; controls lacking one or both of the drivers and effectors were run in key control experiments. Statistics and graphing were carried out with Graphpad Prism, version 10.1.1.

## References

1. Hyman SL, Shores A, North KN. The nature and frequency of cognitive deficits in children with neurofibromatosis type 1. Neurology. 2005;65(7):1037–44. doi: 10.1212/01.wnl.0000179303.72345.ce. PubMed PMID: 16217056.

2. Diggs-Andrews KA, Gutmann DH. Modeling cognitive dysfunction in neurofibromatosis-1. Trends in neurosciences. 2013;36(4):237–47. doi: 10.1016/j.tins.2012.12.002. PubMed PMID: 23312374; PubMed Central PMCID: PMC3622809.

3. Constantino JN, Zhang Y, Holzhauer K, Sant S, Long K, Vallorani A, et al. Distribution and Within-Family Specificity of Quantitative Autistic Traits in Patients with Neurofibromatosis Type I. The Journal of pediatrics. 2015;167(3):621-6 e1. Epub 20150604. doi: 10.1016/j.jpeds.2015.04.075. PubMed PMID: 26051969; PubMed Central PMCID: PMCPMC4792262.

4. Eijk S, Mous SE, Dieleman GC, Dierckx B, Rietman AB, de Nijs PFA, et al. Autism Spectrum Disorder in an Unselected Cohort of Children with Neurofibromatosis Type 1 (NF1). Journal of autism and developmental disorders. 2018;48(7):2278–85. doi: 10.1007/s10803-018-3478-0. PubMed PMID: 29423604; PubMed Central PMCID: PMCPMC5995999.

5. Garg S, Plasschaert E, Descheemaeker MJ, Huson S, Borghgraef M, Vogels A, et al. Autism spectrum disorder profile in neurofibromatosis type I. Journal of autism and developmental disorders. 2015;45(6):1649–57. doi: 10.1007/s10803-014-2321-5. PubMed PMID: 25475362.

6. Hyman SL, Arthur Shores E, North KN. Learning disabilities in children with neurofibromatosis type 1: subtypes, cognitive profile, and attention-deficit-hyperactivity disorder. Developmental medicine and child neurology. 2006;48(12):973–7. doi: 10.1017/S0012162206002131. PubMed PMID: 17109785.

7. Morris SM, Acosta MT, Garg S, Green J, Huson S, Legius E, et al. Disease Burden and Symptom Structure of Autism in Neurofibromatosis Type 1: A Study of the International NF1-ASD Consortium Team (INFACT). JAMA Psychiatry. 2016;73(12):1276–84. doi: 10.1001/jamapsychiatry.2016.2600. PubMed PMID: 27760236; PubMed Central PMCID: PMCPMC5298203.

8. Plasschaert E, Van Eylen L, Descheemaeker MJ, Noens I, Legius E, Steyaert J. Executive functioning deficits in children with neurofibromatosis type 1: The influence of intellectual and social functioning. Am J Med Genet B Neuropsychiatr Genet. 2016;171B(3):348-62. Epub 20160115. doi: 10.1002/ajmg.b.32414. PubMed PMID: 26773288.

9. Walsh KS, Velez JI, Kardel PG, Imas DM, Muenke M, Packer RJ, et al. Symptomatology of autism spectrum disorder in a population with neurofibromatosis type 1. Developmental medicine and child neurology. 2013;55(2):131–8. doi: 10.1111/dmcn.12038. PubMed PMID: 23163951.

10. Vassallo G, Mughal Z, Robinson L, Weisberg D, Roberts SA, Hupton E, et al. Perceived fatigue in children and young adults with neurofibromatosis type 1. J Paediatr Child Health. 2020;56(6):878-83. Epub 20200109. doi: 10.1111/jpc.14764. PubMed PMID: 31916647.

11. Botero V, Stanhope BA, Brown EB, Grenci EC, Boto T, Park SJ, et al. Neurofibromin regulates metabolic rate via neuronal mechanisms in Drosophila. Nature communications. 2021;12(1):4285. Epub 20210713. doi: 10.1038/s41467-021-24505-x. PubMed PMID: 34257279; PubMed Central PMCID: PMCPMC8277851.

12. Botero V, Tomchik SM. Unraveling neuronal and metabolic alterations in neurofibromatosis type 1. J Neurodev Disord. 2024;16(1):49. Epub 20240831. doi: 10.1186/s11689-024-09565-6. PubMed PMID: 39217323; PubMed Central PMCID: PMCPMC11365184.

13. Martin GA, Viskochil D, Bollag G, McCabe PC, Crosier WJ, Haubruck H, et al. The GAP-related domain of the neurofibromatosis type 1 gene product interacts with ras p21. Cell. 1990;63(4):843-9. PubMed PMID: 2121370.

14. Shen MH, Harper PS, Upadhyaya M. Molecular genetics of neurofibromatosis type 1 (NF1). J Med Genet. 1996;33(1):2–17. doi: 10.1136/jmg.33.1.2. PubMed PMID: 8825042; PubMed Central PMCID: PMCPMC1051805.

15. Walker JA, Gouzi JY, Long JB, Huang S, Maher RC, Xia H, et al. Genetic and functional studies implicate synaptic overgrowth and ring gland cAMP/PKA signaling defects in the Drosophila melanogaster neurofibromatosis-1 growth deficiency. PLoS genetics. 2013;9(11):e1003958. doi: 10.1371/journal.pgen.1003958. PubMed PMID: 24278035; PubMed Central PMCID: PMC3836801.

16. Dasgupta B, Dugan LL, Gutmann DH. The neurofibromatosis 1 gene product neurofibromin regulates pituitary adenylate cyclase-activating polypeptide-mediated signaling in astrocytes. The Journal of neuroscience : the official journal of the Society for Neuroscience. 2003;23(26):8949-54. PubMed PMID: 14523097.

17. Tong J, Hannan F, Zhu Y, Bernards A, Zhong Y. Neurofibromin regulates G protein-stimulated adenylyl cyclase activity. Nature neuroscience. 2002;5(2):95–6. doi: 10.1038/nn792. PubMed PMID: 11788835.

18. The I, Hannigan GE, Cowley GS, Reginald S, Zhong Y, Gusella JF, et al. Rescue of a Drosophila NF1 mutant phenotype by protein kinase A. Science. 1997;276(5313):791-4. PubMed PMID: 9115203.

19. Wolman MA, de Groh ED, McBride SM, Jongens TA, Granato M, Epstein JA. Modulation of cAMP and ras signaling pathways improves distinct behavioral deficits in a zebrafish model of neurofibromatosis type 1. Cell reports. 2014;8(5):1265–70. doi: 10.1016/j.celrep.2014.07.054. PubMed PMID: 25176649.

20. Buchanan ME, Davis RL. A distinct set of Drosophila brain neurons required for neurofibromatosis type 1-dependent learning and memory. The Journal of neuroscience : the official journal of the Society for Neuroscience. 2010;30(30):10135–43. doi: 10.1523/JNEUROSCI.0283-10.2010. PubMed PMID: 20668197; PubMed Central PMCID: PMC2917756.

21. Gross AM, Wolters PL, Dombi E, Baldwin A, Whitcomb P, Fisher MJ, et al. Selumetinib in Children with Inoperable Plexiform Neurofibromas. New England Journal of Medicine. 2020;382(15):1430–42. doi: 10.1056/NEJMoa1912735.

22. Moertel CL, Hirbe AC, Shuhaiber HH, Bielamowicz K, Sidhu A, Viskochil D, et al. ReNeu: A Pivotal, Phase IIb Trial of Mirdametinib in Adults and Children With Symptomatic Neurofibromatosis Type 1-Associated Plexiform Neurofibroma. Journal of Clinical Oncology. 2025;43(6):716–29. doi: 10.1200/jco.24.01034. PubMed PMID: 39514826.

23. Abdel-Rahman O, ElHalawani H, Ahmed H. Risk of Selected Cardiovascular Toxicities in Patients With Cancer Treated With MEK Inhibitors: A Comparative Systematic Review and Meta-Analysis. J Glob Oncol. 2015;1(2):73-82. Epub 20151125. doi: 10.1200/JGO.2015.000802. PubMed PMID: 28804776; PubMed Central PMCID: PMCPMC5539872.

24. Avery RA, Trimboli-Heidler C, Kilburn LB. Separation of outer retinal layers secondary to selumetinib. J AAPOS. 2016;20(3):268-71. Epub 20160421. doi: 10.1016/j.jaapos.2016.01.012. PubMed PMID: 27108842; PubMed Central PMCID: PMCPMC4912405.

25. Baldo F, Magnolato A, Barbi E, Bruno I. Selumetinib side effects in children treated for plexiform neurofibromas: first case reports of peripheral edema and hair color change. BMC Pediatr. 2021;21(1):67. Epub 20210206. doi: 10.1186/s12887-021-02530-5. PubMed PMID: 33549085; PubMed Central PMCID: PMCPMC7866429.

26. Chisholm AK, Haebich KM, Pride NA, Walsh KS, Lami F, Ure A, et al. Delineating the autistic phenotype in children with neurofibromatosis type 1. Mol Autism. 2022;13(1):3. Epub 20220104. doi: 10.1186/s13229-02100481-3. PubMed PMID: 34983638; PubMed Central PMCID: PMCPMC8729013.

27. Gutmann DH, Anastasaki C, Gupta A, Hou Y, Morris SM, Payne JM, et al. Cognition and behavior in neurofibromatosis type 1: report and perspective from the Cognition and Behavior in NF1 (CABIN) Task Force. Genes & development. 2025;39(9-10):541-54. Epub 20250502. doi: 10.1101/gad.352629.125. PubMed PMID: 40127956; PubMed Central PMCID: PMCPMC12047663.

28. Robinson JE, Coughlin GM, Hori AM, Cho JR, Mackey ED, Turan Z, et al. Optical dopamine monitoring with dLight1 reveals mesolimbic phenotypes in a mouse model of neurofibromatosis type 1. eLife. 2019;8. Epub 20190923. doi: 10.7554/eLife.48983. PubMed PMID: 31545171; PubMed Central PMCID: PMCPMC6819083.

29. Brown JA, Emnett RJ, White CR, Yuede CM, Conyers SB, O’Malley KL, et al. Reduced striatal dopamine underlies the attention system dysfunction in neurofibromatosis-1 mutant mice. Human molecular genetics. 2010;19(22):4515–28. Epub 20100907. doi: 10.1093/hmg/ddq382. PubMed PMID: 20826448; PubMed Central PMCID: PMCPMC2957316.

30. Sullivan K, El-Hoss J, Quinlan KG, Deo N, Garton F, Seto JT, et al. NF1 is a critical regulator of muscle development and metabolism. Human molecular genetics. 2014;23(5):1250–9. Epub 20131024. doi: 10.1093/hmg/ddt515. PubMed PMID: 24163128; PubMed Central PMCID: PMCPMC3954124.

31. Masgras I, Ciscato F, Brunati AM, Tibaldi E, Indraccolo S, Curtarello M, et al. Absence of Neurofibromin Induces an Oncogenic Metabolic Switch via Mitochondrial ERK-Mediated Phosphorylation of the Chaperone TRAP1. Cell Reports. 2017;18(3):659–72. doi: 10.1016/j.celrep.2016.12.056.

32. Souza MLR, Jansen AK, Rodrigues LOC, Vilela DLS, Kakehasi AM, Martins AS, et al. Increased resting metabolism in neurofibromatosis type 1. Clin Nutr ESPEN. 2019;32:44–9. Epub 20190524. doi: 10.1016/j.clnesp.2019.05.006. PubMed PMID: 31221289.

33. Apostolova I, Derlin T, Salamon J, Amthauer H, Granstrom S, Brenner W, et al. Cerebral glucose metabolism in adults with neurofibromatosis type 1. Brain Res. 2015;1625:97–101. Epub 20150901. doi: 10.1016/j.brainres.2015.08.025. PubMed PMID: 26335059.

34. Martins AS, Jansen AK, Rodrigues LO, Matos CM, Souza ML, de Souza JF, et al. Lower fasting blood glucose in neurofibromatosis type 1. Endocr Connect. 2016;5(1):28–33. doi: 10.1530/EC-15-0102. PubMed PMID: 26631381; PubMed Central PMCID: PMCPMC4703950.

35. Souza M, Jansen A, Martins A, Rodrigues L, Rezende N. Body composition in adults with neurofibromatosis type 1. Rev Assoc Med Bras (1992). 2016;62(9):831–6. doi: 10.1590/1806-9282.62.09.831. PubMed PMID: 28001256.

36. Souza JF, Passos RL, Guedes AC, Rezende NA, Rodrigues LO. Muscular force is reduced in neurofibromatosis type 1. J Musculoskelet Neuronal Interact. 2009;9(1):15-7. PubMed PMID: 19240363.

37. Tritz R, Benson T, Harris V, Hudson FZ, Mintz J, Zhang H, et al. Nf1 heterozygous mice recapitulate the anthropometric and metabolic features of human neurofibromatosis type 1. Transl Res. 2021;228:52–63. Epub 20200808. doi: 10.1016/j.trsl.2020.08.001. PubMed PMID: 32781282; PubMed Central PMCID: PMCPMC7779711.

38. Wei X, Franke J, Ost M, Wardelmann K, Borno S, Timmermann B, et al. Cell autonomous requirement of neurofibromin (Nf1) for postnatal muscle hypertrophic growth and metabolic homeostasis. J Cachexia Sarcopenia Muscle. 2020;11(6):1758–78. Epub 20201019. doi: 10.1002/jcsm.12632. PubMed PMID: 33078583; PubMed Central PMCID: PMCPMC7749575.

39. Kossler N, Stricker S, Rodelsperger C, Robinson PN, Kim J, Dietrich C, et al. Neurofibromin (Nf1) is required for skeletal muscle development. Human molecular genetics. 2011;20(14):2697–709. Epub 20110409. doi: 10.1093/hmg/ddr149. PubMed PMID: 21478499; PubMed Central PMCID: PMCPMC3118757.

40. Maurer GW, Malita A, Nagy S, Koyama T, Werge TM, Halberg KA, et al. Analysis of genes within the schizophrenia-linked 22q11.2 deletion identifies interaction of night owl/LZTR1 and NF1 in GABAergic sleep control. PLoS genetics. 2020;16(4):e1008727. Epub 20200427. doi: 10.1371/journal.pgen.1008727. PubMed PMID: 32339168; PubMed Central PMCID: PMCPMC7205319.

41. Tong JJ, Schriner SE, McCleary D, Day BJ, Wallace DC. Life extension through neurofibromin mitochondrial regulation and antioxidant therapy for neurofibromatosis-1 in Drosophila melanogaster. Nature genetics. 2007;39(4):476–85. Epub 20070318. doi: 10.1038/ng2004. PubMed PMID: 17369827.

42. van der Voet M, Harich B, Franke B, Schenck A. ADHD-associated dopamine transporter, latrophilin and neurofibromin share a dopamine-related locomotor signature in Drosophila. Molecular psychiatry. 2015. doi: 10.1038/mp.2015.55. PubMed PMID: 25962619.

43. King LB, Boto T, Botero V, Aviles AM, Jomsky BM, Joseph C, et al. Developmental loss of neurofibromin across distributed neuronal circuits drives excessive grooming in Drosophila. PLoS genetics. 2020;16(7):e1008920. Epub 20200722. doi: 10.1371/journal.pgen.1008920. PubMed PMID: 32697780; PubMed Central PMCID: PMCPMC7398555.

44. King LB, Koch M, Murphy KR, Velazquez Y, Ja WW, Tomchik SM. Neurofibromin Loss of Function Drives Excessive Grooming in Drosophila. G3 (Bethesda). 2016;6(4):1083–93. Epub 20160407. doi: 10.1534/g3.115.026484. PubMed PMID: 26896440; PubMed Central PMCID: PMCPMC4825643.

45. Suarez GO, Kumar DS, Brunner H, Knauss A, Barrios J, Emel J, et al. Neurofibromin Deficiency Alters the Patterning and Prioritization of Motor Behaviors in a State-Dependent Manner. The Journal of Neuroscience. 2025;45(16):e1531242025. doi: 10.1523/jneurosci.1531-24.2025.

46. Brown EB, Zhang J, Lloyd E, Lanzon E, Botero V, Tomchik S, et al. Neurofibromin 1 mediates sleep depth in Drosophila. PLoS genetics. 2023;19(12):e1011049. Epub 20231213. doi: 10.1371/journal.pgen.1011049. PubMed PMID: 38091360; PubMed Central PMCID: PMCPMC10763969.

47. Vanhaesebroeck B, Guillermet-Guibert J, Graupera M, Bilanges B. The emerging mechanisms of isoform-specific PI3K signalling. Nat Rev Mol Cell Biol. 2010;11(5):329–41. Epub 20100409. doi: 10.1038/nrm2882. PubMed PMID: 20379207.

48. Chowanski S, Walkowiak-Nowicka K, Winkiel M, Marciniak P, Urbanski A, Pacholska-Bogalska J. Insulin-Like Peptides and Cross-Talk With Other Factors in the Regulation of Insect Metabolism. Front Physiol. 2021;12:701203. Epub 20210629. doi: 10.3389/fphys.2021.701203. PubMed PMID: 34267679; PubMed Central PMCID: PMCPMC8276055.

49. Williams JA, Su HS, Bernards A, Field J, Sehgal A. A circadian output in Drosophila mediated by neurofibromatosis-1 and Ras/MAPK. Science. 2001;293(5538):2251–6. doi: 10.1126/science.1063097. PubMed PMID: 11567138.

50. Gouzi JY, Moressis A, Walker JA, Apostolopoulou AA, Palmer RH, Bernards A, et al. The receptor tyrosine kinase Alk controls neurofibromin functions in Drosophila growth and learning. PLoS genetics. 2011;7(9):e1002281. doi: 10.1371/journal.pgen.1002281. PubMed PMID: 21949657; PubMed Central PMCID: PMC3174217.

51. Guo HF, The I, Hannan F, Bernards A, Zhong Y. Requirement of Drosophila NF1 for activation of adenylyl cyclase by PACAP38-like neuropeptides. Science. 1997;276(5313):795-8. PubMed PMID: 9115204.

52. Dyson A, Ryan M, Garg S, Evans DG, Baines RA. Loss of NF1 in Drosophila larvae causes tactile hypersensitivity and impaired synaptic transmission at the neuromuscular junction. The Journal of neuroscience : the official journal of the Society for Neuroscience. 2022. Epub 20221103. doi: 10.1523/JNEUROSCI.0562-22.2022. PubMed PMID: 36344265.

53. Durkin J, Poe AR, Belfer SJ, Rodriguez A, Tang SH, Walker JA, et al. Neurofibromin 1 regulates early developmental sleep in Drosophila. Neurobiol Sleep Circadian Rhythms. 2023;15:100101. Epub 20230809. doi: 10.1016/j.nbscr.2023.100101. PubMed PMID: 37593040; PubMed Central PMCID: PMCPMC10428071.

54. Brown EB, Zhang J, Lloyd E, Lanzon E, Botero V, Tomchik SM, et al. Neurofibromin 1 mediates sleep depth in Drosophila. bioRxiv. 2022. doi: doi:10.1101/2022.09.15.508161.

55. Yatsenko AS, Marrone AK, Kucherenko MM, Shcherbata HR. Measurement of metabolic rate in Drosophila using respirometry. J Vis Exp. 2014;(88):e51681. Epub 20140624. doi: 10.3791/51681. PubMed PMID: 24998593; PubMed Central PMCID: PMCPMC4205100.

56. Saxton RA, Sabatini DM. mTOR Signaling in Growth, Metabolism, and Disease. Cell. 2017;169(2):361–71. doi: 10.1016/j.cell.2017.03.035. PubMed PMID: 28388417.

57. Johannessen CM, Johnson BW, Williams SM, Chan AW, Reczek EE, Lynch RC, et al. TORC1 is essential for NF1-associated malignancies. Current biology : CB. 2008;18(1):56–62. Epub 20071227. doi: 10.1016/j.cub.2007.11.066. PubMed PMID: 18164202.

58. Ullrich NJ, Prabhu SP, Reddy AT, Fisher MJ, Packer R, Goldman S, et al. A phase II study of continuous oral mTOR inhibitor everolimus for recurrent, radiographic-progressive neurofibromatosis type 1-associated pediatric low-grade glioma: a Neurofibromatosis Clinical Trials Consortium study. Neuro Oncol. 2020;22(10):1527–35. doi: 10.1093/neuonc/noaa071. PubMed PMID: 32236425; PubMed Central PMCID: PMCPMC7566451.

59. Kaul A, Toonen JA, Cimino PJ, Gianino SM, Gutmann DH. Aktor MEKmediated mTOR inhibition suppresses Nf1 optic glioma growth. Neuro Oncol. 2015;17(6):843–53. Epub 20141221. doi: 10.1093/neuonc/nou329. PubMed PMID: 25534823; PubMed Central PMCID: PMCPMC4483119.

60. Scheidenhelm DK, Cresswell J, Haipek CA, Fleming TP, Mercer RW, Gutmann DH. Akt-dependent cell size regulation by the adhesion molecule on glia occurs independently of phosphatidylinositol 3-kinase and Rheb signaling. Molecular and cellular biology. 2005;25(8):3151–62. doi: 10.1128/MCB.25.8.3151-3162.2005. PubMed PMID: 15798201; PubMed Central PMCID: PMCPMC1069598.

61. He L, Cho S, Blenis J. mTORC1, the maestro of cell metabolism and growth. Genes & development. 2025;39(1-2):109-31. Epub 20250107. doi: 10.1101/gad.352084.124. PubMed PMID: 39572234; PubMed Central PMCID: PMCPMC11789495.

62. Panwar V, Singh A, Bhatt M, Tonk RK, Azizov S, Raza AS, et al. Multifaceted role of mTOR (mammalian target of rapamycin) signaling pathway in human health and disease. Signal Transduct Target Ther. 2023;8(1):375. Epub 20231002. doi: 10.1038/s41392-023-01608-z. PubMed PMID: 37779156; PubMed Central PMCID: PMCPMC10543444.

63. Guo HF, Tong J, Hannan F, Luo L, Zhong Y. A neurofibromatosis-1-regulated pathway is required for learning in Drosophila. Nature. 2000;403(6772):895–8. doi: 10.1038/35002593. PubMed PMID: 10706287.

64. Walker JA, Tchoudakova AV, McKenney PT, Brill S, Wu D, Cowley GS, et al. Reduced growth of Drosophila neurofibromatosis 1 mutants reflects a non-cell-autonomous requirement for GTPase-Activating Protein activity in larval neurons. Genes & development. 2006;20(23):3311–23. doi: 10.1101/gad.1466806. PubMed PMID: 17114577; PubMed Central PMCID: PMC1686607.

65. Machado Almeida P, Lago Solis B, Stickley L, Feidler A, Nagoshi E. Neurofibromin 1 in mushroom body neurons mediates circadian wake drive through activating cAMP-PKA signaling. Nature communications. 2021;12(1):5758. Epub 20211001. doi: 10.1038/s41467-021-26031-2. PubMed PMID: 34599173; PubMed Central PMCID: PMCPMC8486785.

66. Levin LR, Han PL, Hwang PM, Feinstein PG, Davis RL, Reed RR. The Drosophila learning and memory gene rutabaga encodes a Ca2+/Calmodulin-responsive adenylyl cyclase. Cell. 1992;68(3):479–89. doi: 10.1016/0092-8674(92)90185-f. PubMed PMID: 1739965.

67. Mendoza MC, Er EE, Blenis J. The Ras-ERK and PI3K-mTOR pathways: cross-talk and compensation. Trends Biochem Sci. 2011;36(6):320–8. Epub 20110430. doi: 10.1016/j.tibs.2011.03.006. PubMed PMID: 21531565; PubMed Central PMCID: PMCPMC3112285.

68. Gronke S, Beller M, Fellert S, Ramakrishnan H, Jackle H, Kuhnlein RP. Control of fat storage by a Drosophila PAT domain protein. Current biology : CB. 2003;13(7):603–6. doi: 10.1016/s0960-9822(03)00175-1. PubMed PMID: 12676093.

69. Dickerson BH, de Souza AM, Huda A, Dickinson MH. Flies Regulate Wing Motion via Active Control of a Dual-Function Gyroscope. Current Biology. 2019;29(20):3517-24.e3. doi: 10.1016/j.cub.2019.08.065.

70. Musselman LP, Kuhnlein RP. Drosophila as a model to study obesity and metabolic disease. J Exp Biol. 2018;221(Pt Suppl 1). Epub 20180307. doi: 10.1242/jeb.163881. PubMed PMID: 29514880.

71. Huang K, Liu Y, Perrimon N. Roles of Insect Oenocytes in Physiology and Their Relevance to Human Metabolic Diseases. Front Insect Sci. 2022;2:859847. Epub 20220317. doi: 10.3389/finsc.2022.859847. PubMed PMID: 38468774; PubMed Central PMCID: PMCPMC10926422.

72. Takemura S-y, Hayworth KJ, Huang GB, Januszewski M, Lu Z, Marin EC, et al. A Connectome of the Male Drosophila Ventral Nerve Cord. eLife Sciences Publications, Ltd; 2024.

73. Summers MA, Quinlan KG, Payne JM, Little DG, North KN, Schindeler A. Skeletal muscle and motor deficits in Neurofibromatosis Type 1. J Musculoskelet Neuronal Interact. 2015;15(2):161-70. PubMed PMID: 26032208; PubMed Central PMCID: PMCPMC5133719.

74. Chatterjee N, Perrimon N. What fuels the fly: Energy metabolism in Drosophila and its application to the study of obesity and diabetes. Science Advances. 2021;7(24):eabg4336. doi: doi:10.1126/sciadv.abg4336.

75. Wegener G. Flying insects: model systems in exercise physiology. Experientia. 1996;52(5):404–12. doi: 10.1007/bf01919307. PubMed PMID: 8641375.

76. Gutman DH, Andersen LB, Cole JL, Swaroop M, Collins FS. An alternatively-spliced mRNA in the carboxy terminus of the neurofibromatosis type 1 (NF1) gene is expressed in muscle. Human molecular genetics. 1993;2(7):989–92. doi: 10.1093/hmg/2.7.989. PubMed PMID: 8364582.

77. Barwell T, DeVeale B, Poirier L, Zheng J, Seroude F, Seroude L. Regulating the UAS/GAL4 system in adult Drosophila with Tet-off GAL80 transgenes. PeerJ. 2017;5:e4167. Epub 20171214. doi: 10.7717/peerj.4167. PubMed PMID: 29259847; PubMed Central PMCID: PMCPMC5733373.

78. Weaver LN, Ma T, Drummond-Barbosa D. Analysis of Gal4 Expression Patterns in Adult Drosophila Females. G3 Genes|Genomes|Genetics. 2020;10(11):4147–58. doi: 10.1534/g3.120.401676.

79. Gonzalez E, McGraw TE. The Akt kinases: isoform specificity in metabolism and cancer. Cell Cycle. 2009;8(16):2502–8. Epub 20090823. doi: 10.4161/cc.8.16.9335. PubMed PMID: 19597332; PubMed Central PMCID: PMCPMC2997486.

80. Lee G, Chung J. Discrete functions of rictor and raptor in cell growth regulation in Drosophila. Biochem Biophys Res Commun. 2007;357(4):11549. Epub 20070420. doi: 10.1016/j.bbrc.2007.04.086. PubMed PMID: 17462592.

81. Watson AL, Anderson LK, Greeley AD, Keng VW, Rahrmann EP, Halfond AL, et al. Co-targeting the MAPK and PI3K/AKT/mTOR pathways in two genetically engineered mouse models of schwann cell tumors reduces tumor grade and multiplicity. Oncotarget. 2014;5(6):1502–14. doi: 10.18632/oncotarget.1609. PubMed PMID: 24681606; PubMed Central PMCID: PMCPMC4039227.

82. Parra V, Verdejo HE, Iglewski M, Del Campo A, Troncoso R, Jones D, et al. Insulin stimulates mitochondrial fusion and function in cardiomyocytes via the Akt-mTOR-NFκB-Opa-1 signaling pathway. Diabetes. 2014;63(1):75–88. Epub 20130905. doi: 10.2337/db13-0340. PubMed PMID: 24009260; PubMed Central PMCID: PMCPMC3868041.

83. Anastasaki C, Gutmann DH. Neuronal NF1/RAS regulation of cyclic AMP requires atypical PKC activation. Human molecular genetics. 2014;23(25):6712–21. Epub 20140728. doi: 10.1093/hmg/ddu389. PubMed PMID: 25070947; PubMed Central PMCID: PMCPMC4245041.

84. Dasgupta B, Yi Y, Chen DY, Weber JD, Gutmann DH. Proteomic analysis reveals hyperactivation of the mammalian target of rapamycin pathway in neurofibromatosis 1-associated human and mouse brain tumors. Cancer Res. 2005;65(7):2755–60. doi: 10.1158/0008-5472.CAN-04-4058. PubMed PMID: 15805275.

85. Johannessen CM, Reczek EE, James MF, Brems H, Legius E, Cichowski K. The NF1 tumor suppressor critically regulates TSC2 and mTOR. Proceedings of the National Academy of Sciences of the United States of America. 2005;102(24):8573–8. Epub 20050603. doi: 10.1073/pnas.0503224102. PubMed PMID: 15937108; PubMed Central PMCID: PMCPMC1142482.

86. Anastasaki C, Woo AS, Messiaen LM, Gutmann DH. Elucidating the impact of neurofibromatosis-1 germline mutations on neurofibromin function and dopamine-based learning. Human molecular genetics. 2015;24(12):3518–28. Epub 20150318. doi: 10.1093/hmg/ddv103. PubMed PMID: 25788518; PubMed Central PMCID: PMCPMC4553714.

87. Cui Y, Costa RM, Murphy GG, Elgersma Y, Zhu Y, Gutmann DH, et al. Neurofibromin regulation of ERK signaling modulates GABA release and learning. Cell. 2008;135(3):549–60. doi: 10.1016/j.cell.2008.09.060. PubMed PMID: 18984165; PubMed Central PMCID: PMC2673196.

88. Kikuchi A, Demo SD, Ye ZH, Chen YW, Williams LT. ralGDS family members interact with the effector loop of ras p21. Molecular and cellular biology. 1994;14(11):7483–91. doi: 10.1128/mcb.14.11.7483-7491.1994. PubMed PMID: 7935463; PubMed Central PMCID: PMCPMC359284.

89. Bodempudi V, Yamoutpoor F, Pan W, Dudek AZ, Esfandyari T, Piedra M, et al. Ral overactivation in malignant peripheral nerve sheath tumors. Molecular and cellular biology. 2009;29(14):3964–74. Epub 20090504. doi: 10.1128/MCB.01153-08. PubMed PMID: 19414599; PubMed Central PMCID: PMCPMC2704746.

90. Danielsen SA, Lind GE, Kolberg M, Holand M, Bjerkehagen B, Sundby Hall K, et al. Methylated RASSF1A in malignant peripheral nerve sheath tumors identifies neurofibromatosis type 1 patients with inferior prognosis. Neuro Oncol. 2015;17(1):63–9. Epub 20140719. doi: 10.1093/neuonc/nou140. PubMed PMID: 25038505; PubMed Central PMCID: PMCPMC4416132.

91. Mund JA, Park S, Smith AE, He Y, Jiang L, Hawley E, et al. Genetic disruption of the small GTPase RAC1 prevents plexiform neurofibroma formation in mice with neurofibromatosis type 1. J Biol Chem. 2020;295(29):9948–58. Epub 20200529. doi: 10.1074/jbc.RA119.010981. PubMed PMID: 32471868; PubMed Central PMCID: PMCPMC7380178.

92. Kohlmeyer JL, Lingo JJ, Kaemmer CA, Scherer A, Warrier A, Voigt E, et al. CDK4/6-MEK Inhibition in MPNSTs Causes Plasma Cell Infiltration, Sensitization to PD-L1 Blockade, and Tumor Regression. Clin Cancer Res. 2023;29(17):3484–97. doi: 10.1158/1078-0432.CCR-23-0749. PubMed PMID: 37410426; PubMed Central PMCID: PMCPMC10528807.

93. Ortega-Bertran S, Fernandez-Rodriguez J, Magallon-Lorenz M, Zhang X, Creus-Bachiller E, Diazgranados AP, et al. Triple Combination of MEK, BET, and CDK Inhibitors Significantly Reduces Human Malignant Peripheral Nerve Sheath Tumors in Mouse Models. Clin Cancer Res. 2025;31(5):907–20. doi: 10.1158/1078-0432.CCR-24-2807. PubMed PMID: 39786423; PubMed Central PMCID: PMCPMC11873804.

94. Delandre C, McMullen JPD, Marshall OJ. Dynamic changes in neuronal and glial GAL4 driver expression during Drosophila aging. Genetics. 2025;229(3). doi: 10.1093/genetics/iyaf014. PubMed PMID: 39950543; PubMed Central PMCID: PMCPMC11912842.

95. Weaver LN, Ma T, Drummond-Barbosa D. Analysis of Gal4 Expression Patterns in Adult Drosophila Females. G3 (Bethesda). 2020;10(11):414758. Epub 20201105. doi: 10.1534/g3.120.401676. PubMed PMID: 32917721; PubMed Central PMCID: PMCPMC7642949.

96. Sofela FA, Lopez Valencia M, Jongens TA, Sehgal A. Effects of Nf1 on sleep behavior are mediated through starvation caused by deficits in SARM1 dependent NAD+ metabolism. bioRxiv. 2024:2024.09.14.612058. doi: 10.1101/2024.09.14.612058.

97. Dietzl G, Chen D, Schnorrer F, Su KC, Barinova Y, Fellner M, et al. A genome-wide transgenic RNAi library for conditional gene inactivation in Drosophila. Nature. 2007;448(7150):151–6. doi: 10.1038/nature05954. PubMed PMID: 17625558.

98. Szebenyi SJ. Cleaning behaviour in Drosophila melanogaster. Animal behaviour. 1969;17(4):641–51.

99. Jenett A, Rubin GM, Ngo TT, Shepherd D, Murphy C, Dionne H, et al. A GAL4-driver line resource for Drosophila neurobiology. Cell reports. 2012;2(4):991–1001. Epub 20121011. doi: 10.1016/j.celrep.2012.09.011. PubMed PMID: 23063364; PubMed Central PMCID: PMCPMC3515021.

